# Interdependence of intra- and inter-domain motions in the PSD-95 PDZ12 tandem

**DOI:** 10.1101/640631

**Authors:** Bertalan Kovács, Nóra Zajácz-Epresi, Zoltán Gáspári

## Abstract

PSD-95 is the most abundant scaffold protein in the postsynaptic density of neurons. Its two N-terminal PDZ domains form an autonomous structural unit and their interdomain orientation and dynamics was shown to be dependent on binding to various partner proteins. To understand the mechanistic details of the effect of ligand binding on interdomain structure and dynamics, we generated conformational ensembles using experimentally determined NOE interatomic distances and S^2^ order parameters, available from the literature. In our approach no explicit restraints between the two domains were used and their fast dynamics was also treated independently. We found that intradomain structural changes induced by ligand binding have a profound effect on the interfaces where interdomain contacts can be formed, modulating the probability of the occurrence of specific domain-domain orientations. Our results suggest that the β2-β3 loop in the PDZ domains is a key regulatory region that, through interacting with the upstream residues of the C-terminal peptide ligand, influences both intradomain motions and supramodular rearrangement.

## Introduction

PDZ domains (**P**SD-95/**D**iscs-large/**Z**O1), also known as DHR (**d**iscs-large **h**omology **r**egion) or GLGF repeats for their conserved ligand binding motif, are common protein-binding modules with about 270 copies occurring in 150 proteins in the human proteome (Ernst et al., 2014; Luck et al., 2012). While being involved in maintaining epithelial cell polarity, regulating signaling pathways and establishing tight junctions in metazoans, their most widely studied role is organizing synaptic complexes by clustering membrane receptors and ion channels in the postsynaptic density (Fanning and Anderson, 1999; Harris and Lim, 2001; Hung and Sheng, 2002; Luck et al., 2012; Nourry et al., 2003; Sheng and Sala, 2001).

The PDZ domain is often regarded the glue that binds signaling complexes together in nerve cells. The PSD-95 (**p**ost**s**ynaptic **d**ensity protein 95, also referred to as SAP-90 or DLG4 by its encoding gene in humans), member of the MAGUK family (**m**embrane **a**ssociated **gu**anylate **k**inase), contains multiple PDZ domains, which allows it to function not only as a scaffold, but as an adaptor protein to downstream intracellular signaling pathways (Craven and Bredt, 1998; Feng and Zhang, 2009; Harris and Lim, 2001; Kim and Sheng, 2004; Sheng and Sala, 2001). Its abundance in the postsynaptic density and a number of interaction partners have made it a frequent research subject to understand signal transduction (Kim and Sheng, 2004). Because of its role in regulating synaptic plasticity, PSD-95 is implicated in several neurological disorders, like Alzheimer’s disease or neuropathic pain, and its PDZ domains are favored drug targets (Gardoni et al., 2009; Kim and Sheng, 2004).

The approximately 90 residue-long PDZ domains contain 6 β-strands in a semi-open barrel-like formation, which is flanked by 2 α-helices (Figure 1.) (Doyle et al., 1996; Morais Cabral et al., 1996). The ligand binding site recognizes short C-terminal peptide sequences, with the C-terminal hydrophobic residue being accommodated at the carboxylate-binding loop (Harris and Lim, 2001; Nourry et al., 2003). The traditional categorization of PDZ-binding motifs into three classes based solely on the –1 residue of the ligand (Harris and Lim, 2001; Hung and Sheng, 2002) does not explain the subtle differences in the binding affinities. Lately, more complicated specificity trees were introduced and a growing number of evidence indicate the importance of the poorly conserved β2-β3 loop interacting with upstream ligand residues (Ernst et al., 2014; Imamura et al., 2002; Luck et al., 2012; Mostarda et al., 2012; Tochio et al., 2000; Tonikian et al., 2008).

**Figure 1:**
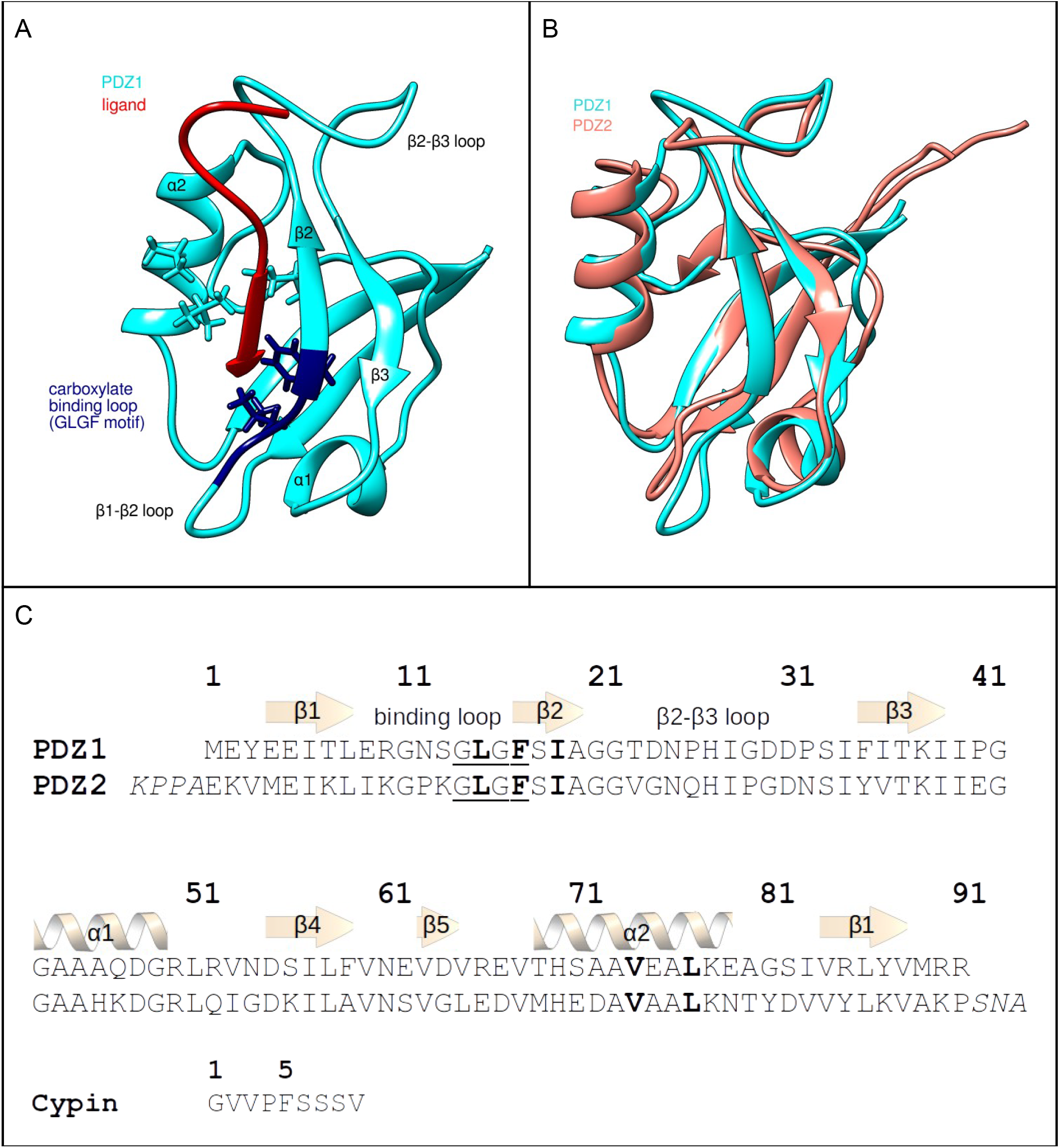
The PDZ12 tandem of PSD95, taken from the PDB structure 2KA9. **A.** Structure of PDZ1 bound to the 9 C-terminal residues of cypin (red). Atoms of the hydrophobic core forming residues are explicitly shown, and the carboxylate binding loop (GLGF motif) is colored dark blue. **B.** Superimposed structures of PDZ1 (cyan) and PDZ2 (salmon). **C.** Sequence alignment of PDZ1 and PDZ2 domains and the sequence of the ligand: 9 C-terminal residues of cypin. Residues of the carboxylate binding loop (GLGF motif) are underlined, residues forming the hydrophobic core are highlighted in bold whereas the linker region preceding and tail the region following the PDZ2 domain is italicized.

NMR studies revealed that internal dynamics is an inherent property of PDZ domains. In PDZ3 of PSD-95, an extra C-terminal helix was shown to regulate ligand-dependent side-chain reorientation (Mostarda et al., 2012; Petit et al., 2009). In PDZ2 of the human protein tyrosine phosphatase 1E (hPTP1E), side-chain order parameters revealed a network that triggers changes in the dynamic behaviour far from the binding site (Dhulesia et al., 2008; Fuentes et al., 2004, 2006).

Neighbouring PDZ domains often form structural and functional supramodules termed PDZ tandems (Feng and Zhang, 2009; Wang et al., 2010; Ye and Zhang, 2013). Such supramodules usually exhibit a biological function different from the simple sum of the two isolated domains. The strength and nature of the interaction between the two domains can vary on a wide range. The PDZ12 tandem of human syntenin adopts a rigid structure, and its activity is regulated by an N-terminal tail region (Cierpicki et al., 2005; Kang et al., 2003). Similarly, in the PDZ12 tandem of X11/Mint proteins, the C-terminal tail has an auto-inhibitory and regulatory role (Long et al., 2005). In turn, both PDZ12 and PDZ45 tandems of GRIP1 exhibit strong interdependence: the sole known function of one of the PDZ domains of the tandem is an intramolecular chaperoning effect, without which the other PDZ domain remains unfolded (Feng et al., 2003; Long et al., 2008; Zhang et al., 2001).

The supramoldular structure and dynamics of PSD-95 has been of great interest, and is suggested to be a key feature which might even influence how it selects amongst its many interaction partners (McCann et al., 2011). However, the exact role and the mode of modulation of interdomain flexibility of its PDZ12 tandem remains elusive. The first crystal structure of PSD-95 PDZ12 tandem showed a rigid interdomain orientation with the ligands pointing to the same direction (Long et al., 2003). Such fixed interdomain orientation could allow for synergistic binding of membrane receptors in the postsynaptic density. Later, however, the supramodular structure of PDZ12 tandem determined by FRET measurements indicated a different interdomain orientation in which the peptide binding grooves pointed to the opposite direction (McCann et al., 2012, 2011). Also, this study came to the conclusion that the same supramodular structure is maintained in the full-length protein as in the isolated tandem (McCann et al., 2011). The FRET structure was subsequently refined with discrete molecular dynamic simulation and was found to be the average of many possible interdomain orientations, with two structures – an open-like and a closed-like conformation – emerging as the energetically most favourable ones (Yanez Orozco et al., 2018). This study also identified the ultra-weak interdomain interactions in the two most dominant conformations that stabilize the supramodular structure of the PDZ12 tandem. Furthermore, it was shown by solution NMR experiments that the interdomain orientation is ligand dependent: the free form of PDZ12 tandem is rigid, whereas ligand binding induces interdomain mobility (Wang et al., 2009).

In this study, to elucidate the supramodular dynamics in the PDZ tandem and the exact atomic-level mechanism of the ligand-induced interdomain allostery, dynamic structural ensembles were generated incorporating NMR-derived distance restraints and backbone order parameters. Our results suggest interdependence of intra- and interdomain motions and provide atomic-level details of the underlying mechanism.

## Results and discussion

### The generated structural ensembles correspond to the experimental data

To uncover the effect of ligand binding on the internal motions, three dynamic structural ensembles were generated with externally restrained molecular dynamic simulation of the PDZ12 tandem: one of its free state and two of its complex state. In the complex states, both PDZ domains were bound with to the 9 C-terminal residues of the cypin (Firestein et al., 1999; Wang et al., 2009). NOE ^1^H-^1^H distance restraints were applied to all three ensembles, and one of the complex structures was further restrained with S^2^ order parameters. (For the details on generating dynamic structural ensembles with NOE and S^2^ restraints, see Methods.)

Table 1 summarizes the RMSDs of the individual PDZ domains in all ensembles and the correlations between the experimental and back-calculated NMR parameters. Backbone RMSDs indicate that both domains in all ensembles have a well folded, compact structure. Local RMSD values are slightly above average for the N- and C-terminal tails as well as for the β1-β2 and β2-β3 loops (Supplementary Figure 1). Since there are less experimental long-range distance restraints per residue in these regions, this difference in the local RMSD does not necessarily indicate a larger local fluctuation. By applying S^2^ order parameters as restraints, a high correlation can be achieved without compromising chemical shift correlation (Table 1).

**Table 1:** Summary of the RMSD of the original and generated ensembles and their compliance with the experimental parameters. The original ensemble was accessed on the Protein Data Bank under ID 2KA9, the rest were generated by externally restrained MD simulations (see Methods.) Pearson’s correlation coefficient was determined between the experimental and back-calculated parameters. RMSDs were calculated, after superimposition, for the N, C and Cɑ2 helix. These motions mainly differ in the backbone atoms in residues 1-91 (PDZ1 domain) and 96-186 (PDZ2 domain).

| Ensemble | No. of conformers | Chemical shift correl. |  | S<sup>2</sup> correl. | PDZ1 RMSD | PDZ2 RMSD |
| --- | --- | --- | --- | --- | --- | --- |
|  |  | N | H |  |  |  |
| *Original PDB* | 20 | 0.83 | 0.41 | 0.10 | 0.57 | 0.62 |
| *Free* | 1448 | 0.85 | 0.55 | 0.22 | 0.73 | 0.66 |
| *Complex* | 1448 | 0.85 | 0.55 | 0.16 | 0.75 | 0.58 |
| *Complex (S<sup>2</sup> restrained)* | 1448 | 0.85 | 0.55 | 0.88 | 0.67 | 0.63 |

### Dominant motions in the free form are redistributed by ligand binding in a domain specific way

Principal component analysis (PCA) was carried out on the core region of both PDZ domains to uncover the nature of the intradomain motions (for details, see Methods). In PDZ1 domain, most of the largest principal components represent the bending of the flexible β1-β2 and β2-β3 loops, and the displacement of the ɑ2 helix. These motions mainly differ in the 2 helix. These motions mainly differ in the direction of the relative displacement of the flexible parts. In PC1 for example, the ɑ2 helix. These motions mainly differ in the2 helix is moving towards and away from the β1 strand which contains the GLGF-motif characteristic of PDZ domains. This motion corresponds to the opening and closing of the hydrophobic core and ligand binding site (Figure 2.B). The binding site openness can be quantified by measuring the distance between the Cɑ2 helix. These motions mainly differ in the atoms of the two hydrophobic residues LEU77 and PHE17 in PDZ1, (and, equivalently, LEU172 and PHE112 in PDZ2), located on the two opposing sites of the ligand binding cleft, i.e. in α2 helix. These motions mainly differ in the2 and β1, respectively. These two residues accommodate the side chain of the hydrophobic C-terminal valine of the ligand.

**Figure 2:**
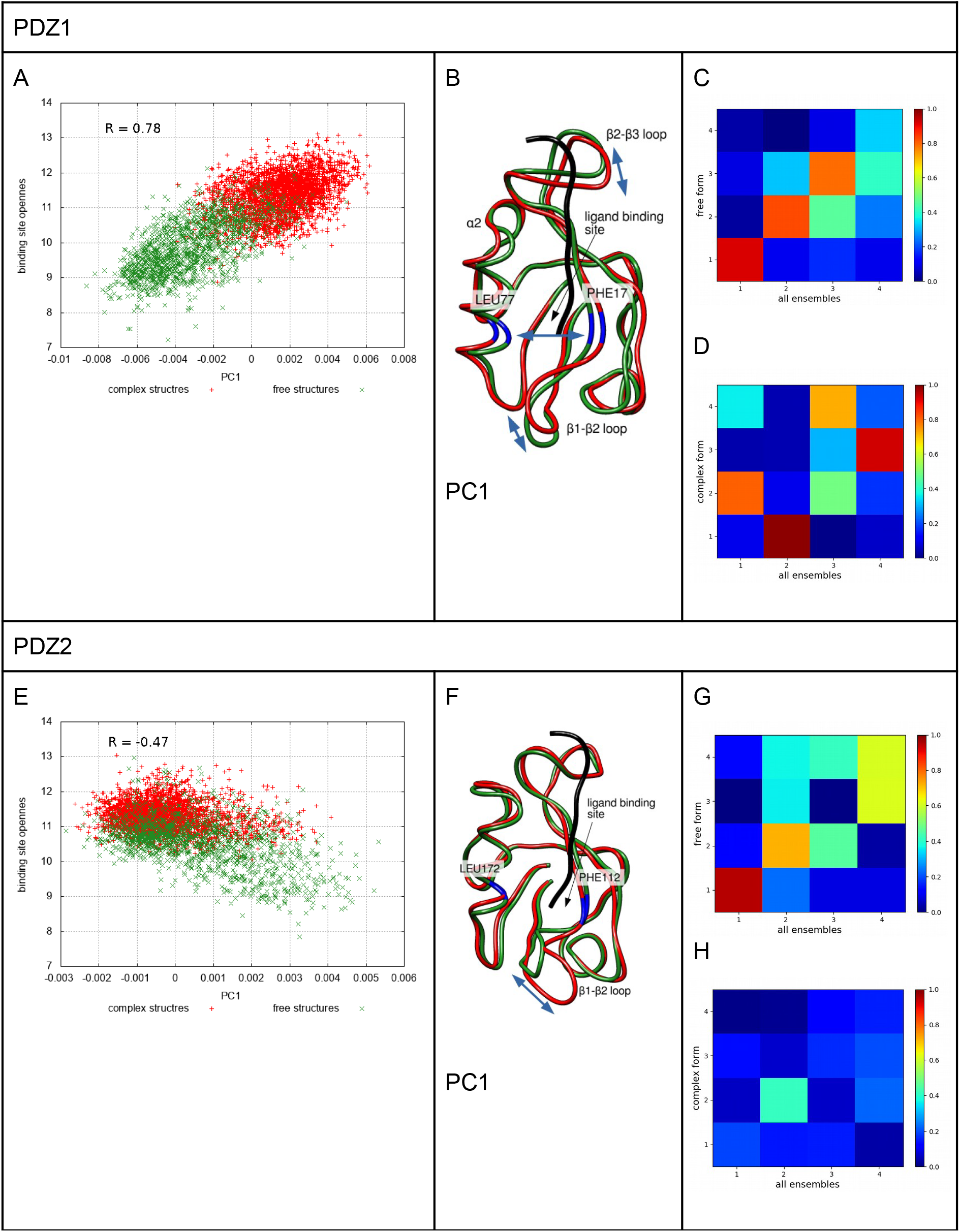
Intradomain motions in PDZ1 (A-D) and PDZ2 (E-H) domains. **A, E**: binding site openness (i.e. distance between Cɑ2 helix. These motions mainly differ in the of PHE17-LEU77 and PHE112-LEU172 in PDZ1 and PDZ2, respectively), plotted against the first principal component (PC1). PC1 for PDZ2 domain in this plot was calculated for the truncated structure (for details, see text). **B, F**: ribbon representation of PC1 in PDZ1 and PDZ2 domains, the most mobile regions are indicated with blue arrow. **C-D,G-H**: overlap between the principal components of the whole ensembles and the free (C,G) and complex (D,H) structures.

The first principal component in PDZ1 correlates well with the binding site openness (R=0.78) (Figure 2.A). The free and complex structures cover a different region along PC1 (and therefore along the binding site openness as well), with the free structures being more extended from the closed binding site almost to the widely open conformation, but the complex structures most frequently occurring in the open conformation. The binding site opening is a motion present both in the free and complex forms of PDZ1, but it is slightly quenched by ligand binding. This is also evidenced by the PC overlap plots (Figures 2.C and 2.D), which indicate that the binding site opening is represented by the first principal component in the free form, but only by the second PC in the complex form. PC1 in the complex form, which shows a high overlap with PC2 in the whole ensemble, represents a shearing displacement of the α2 helix. These motions mainly differ in the 2 helix relative to the β1 strand, as well as the bending of the β1-β2 and β2-β3 loops. However, PC2 in the complex forms does not correlate well with the opening of the binding site (R = 0.39). The high overlap between PC2 in the complex ensembles and PC1 in the whole ensemble is due to the similar motion of of β1-β2 and β2-β3 loops rather than to that of α2 helix. These motions mainly differ in the2 helix, as is evidenced by PCA carried out exclusively on residues of these loops (data not shown). Despite a remarkable increase in S^2^-correlation, no new internal motions in PDZ1 emerge in the S^2^-restrained complex ensemble compared to the unrestrained complex one.

The PDZ2 domain, despite being structurally highly similar to PDZ1, shows some remarkable difference in its internal motions. In the first two principal components the bending of the β1-β2 and β2-β3 loops is dominant, and the α2 helix. These motions mainly differ in the2 helix is but very slightly displaced (Figure 2.F). This results in a low correlation between PC1 or PC2 and the binding site openness (R = 0.41). By excluding the flexible loops (residues 105-109 and 117-127) from the PCA, a slightly better correlation can be achieved (R = –0.47), which is even higher computed for the free structures only (R = –0.50). However, by plotting PC1 of the truncated PDZ2 structure against the binding site openness, the free and the complex structures show some slight separation along both axes (Figure 2.E). The effect, therefore, that the binding of the ligand quenches the binding site opening, is still present in PDZ2, but is far less pronounced than in PDZ1. On the other hand, internal motions in the complex form show poor overlap with those either of the free form or of the whole ensemble. Only the first two PCs of the free form overlap well with those of the whole ensembles (Figures 2.G and 2.H). It can be stated therefore, that in the complex form of PDZ2, new types of internal motions emerge that represent the bending of the β1-β2 loop in PC1, and of the β1-β2 and β2-β3 loops in PC2, but in a different direction.

### Interdomain motions: the relative orientation of the two domains covers a large conformational space

In order to investigate the interdomain mobility, the PDZ1 domains in all 3 ensembles were superimposed to a common template. This results in a very well aligned PDZ1 domain (RMSD = 0.81), whereas the PDZ2 domain samples a large number of different orientations (RMSD = 22.93) highlighting the relative displacement between the two domains. Using these superimposed ensembles, principal component analysis (PCA) was carried out only on the orientations of the core regions of PDZ2 domain relative to PDZ1 (for details, see Methods). The first and second principal components cover 48% and 32% percent of the variability of the structures, respectively.

When plotting the resulting first and second principal components, the data points exhibit a circular arrangement (Figure 3.A). The data points were sampled along two circles that have a radius 0.8 and 1.1 times that of the best fitting circle in a way that the nearest data point was chosen to the 16 equidistant points in both circles (Figure 3.B). Upon visual inspection, it became evident that these data points, which lie along a 2D circle on the PCA plot, represent such PDZ-tandem structures where the PDZ2 domains are arranged along a 3D circle (with the PDZ1 domains being superimposed). Furthermore, the distance of each data point from the center of the circle corresponds to the degree by which the orientation of the two domains deviates from the extended interdomain conformation, i.e. the interdomain bending angle. This observation can be best demonstrated by plotting the centers of mass of the PDZ2 domains that were sampled from the two circles: they fit relatively well to two nearly parallel circles (Figure 3.C).

**Figure 3:**
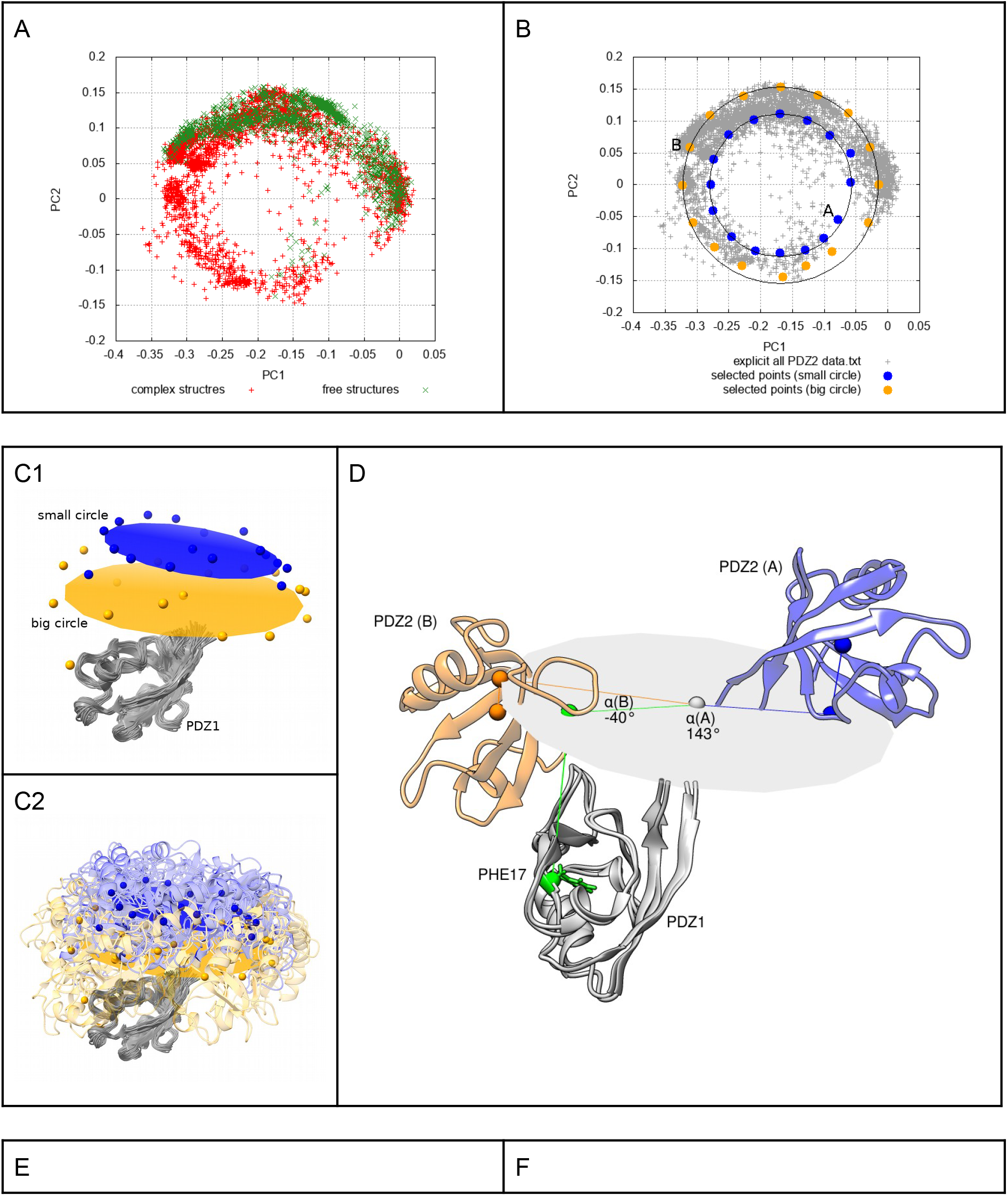
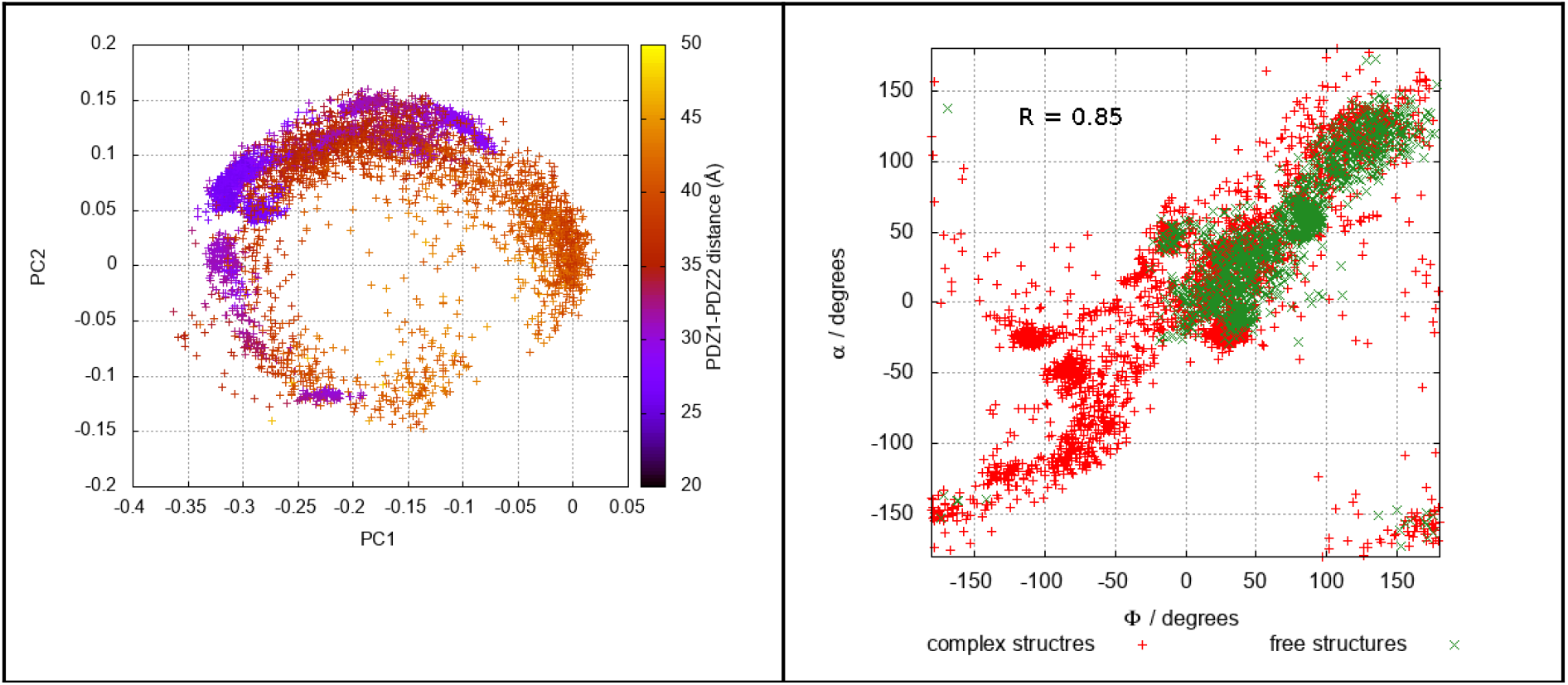
Interdomain motions. **A**: PCA plot of the displacement of PDZ2 domain relative to PDZ1. **B**: two circles fitted to the interdomain PCA plot (with radii 0.8 and 1.1 times of that of the best fitting circle) 2×16 data points are highlighted, which are closest to the 16 equidistant points on each circle. **C1,C2:** The centers of mass of the PDZ2 domain in different orientation relative to PDZ1. **D:** illustration of the interdomain torsion angle (α2 helix. These motions mainly differ in the) in two selected interdomain orientation (also indicated on **B**). For description, see Methods. **E**: PCA plot of the interdomain motions, with color coded distance between the centers of masses of PDZ1 and PDZ2 domains. **F:** Interdomain torsion angle calculated as presented above (α2 helix. These motions mainly differ in the) plotted against the interdomain torsion angle calculated as described by Zhang and coworkers (ϕ) (Wang et al., 2009).

Taking advantage of the exhaustively sampled interdomain conformational space, an interdomain torsion angle (denoted by α) can easily be defined (Figure 3.D, for details, see Methods.) The angle component of the polar coordinates calculated from the first and second principal components, after phase correction, correlate extremely well with the interdomain torsion angle (R = 0.98), and also with the interdomain torsion angle defined by Zhang and coworkers (Wang et al., 2009) (Figure 3.F, denoted ϕ)(R = 0.85).

The presence of the ligand has a remarkable effect on the covered interdomain conformational space: in the complex form, the whole circle is relatively well sampled, whereas in the ligand-free form only a part of the full circle is covered (Figure 3.A). The same effect is also evident from the interdomain torsion angles: the complex form allows for much more extended interdomain motions (Figure 3.F).

Strikingly, the position of the superimposed PDZ1 domain relative to the circle along which PDZ2 domain moves is not perfectly centered, but slightly tilted towards one side (Figure 3.C). This results in a thinly sampled region in the interdomain PCA plot, but – more importantly – a shorter interdomain distance and, consequently, more interdomain contacts on one side of the sampled circle (Figure 3.E).

### Structures forming an interdomain interface can be clustered according to the interdomain orientation

A shorter interdomain distance allows for more interdomain contacts. Structures in which there were at least 20 interdomain atom-atom contacts between heavy atoms within a 5 Å cutoff range (termed “tight structures”) occupy only a fraction of the whole interdomain conformational space (Figure 4.A). Using the polar coordinates, computed with the aid of the best-fitting circle (Figure 3.B), the tight structures can be clustered in 7 clusters by hierarchical clustering algorithm (complete linkage) (Figure 4.A). (For details on clustering the ensembles, see Methods.) The 7 representative structures correspond to 7 tightly-packed interdomain orientations, in which the interdomain interface is formed between different regions (Figure 4.B). For better visibility, the surface of the PDZ domain was divided into 5 sites and termed head, tail, back, left side and right side, relative to the ligand binding site (Figure 4.C, Supplementary Figure 2). Figure 4.D summarizes the interface-forming sites in PDZ1 and PDZ2 domains, as well as the alignment of the two ligands. For more details on the clusters, see Supplementary Table 1.

**Figure 4:**
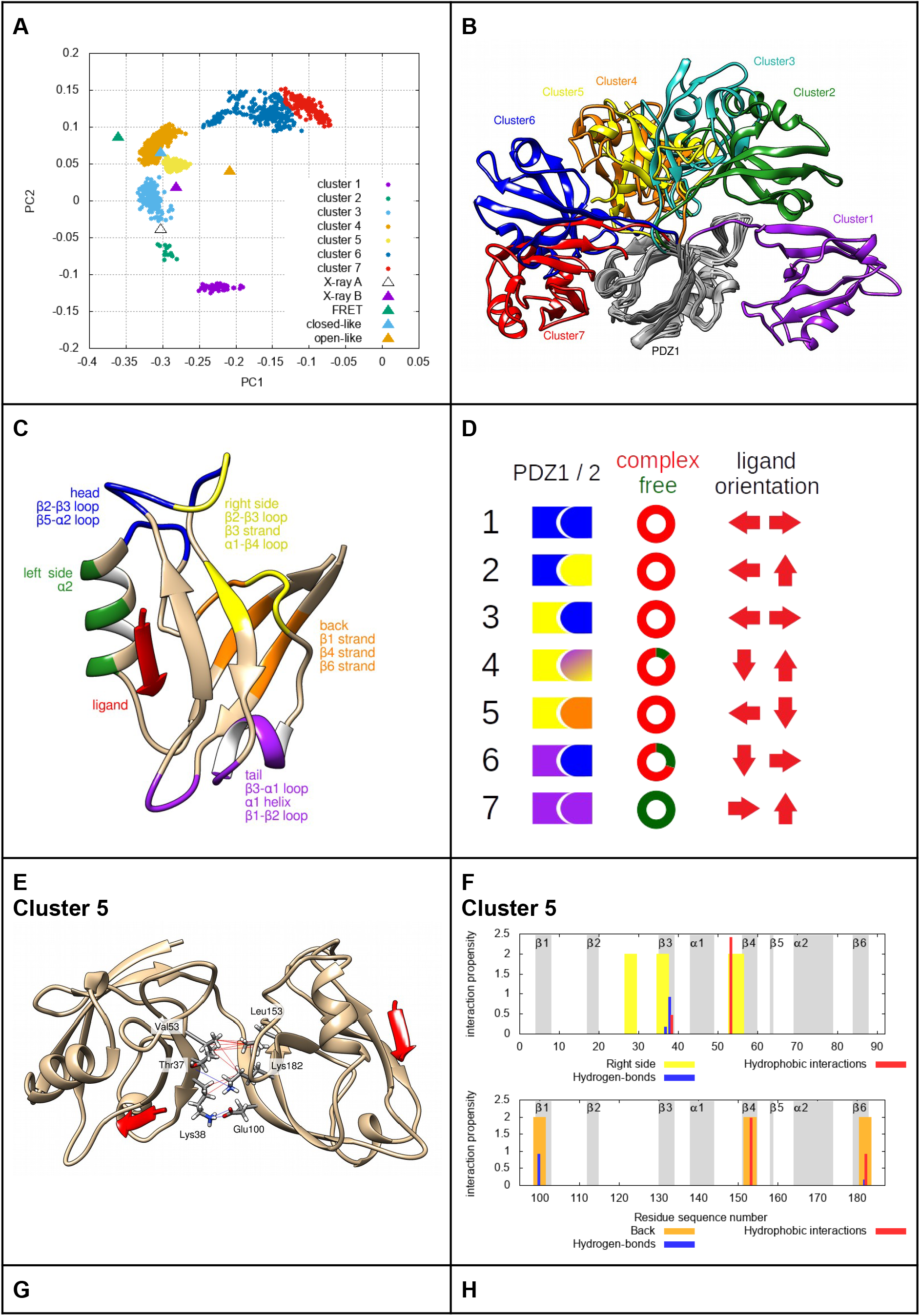
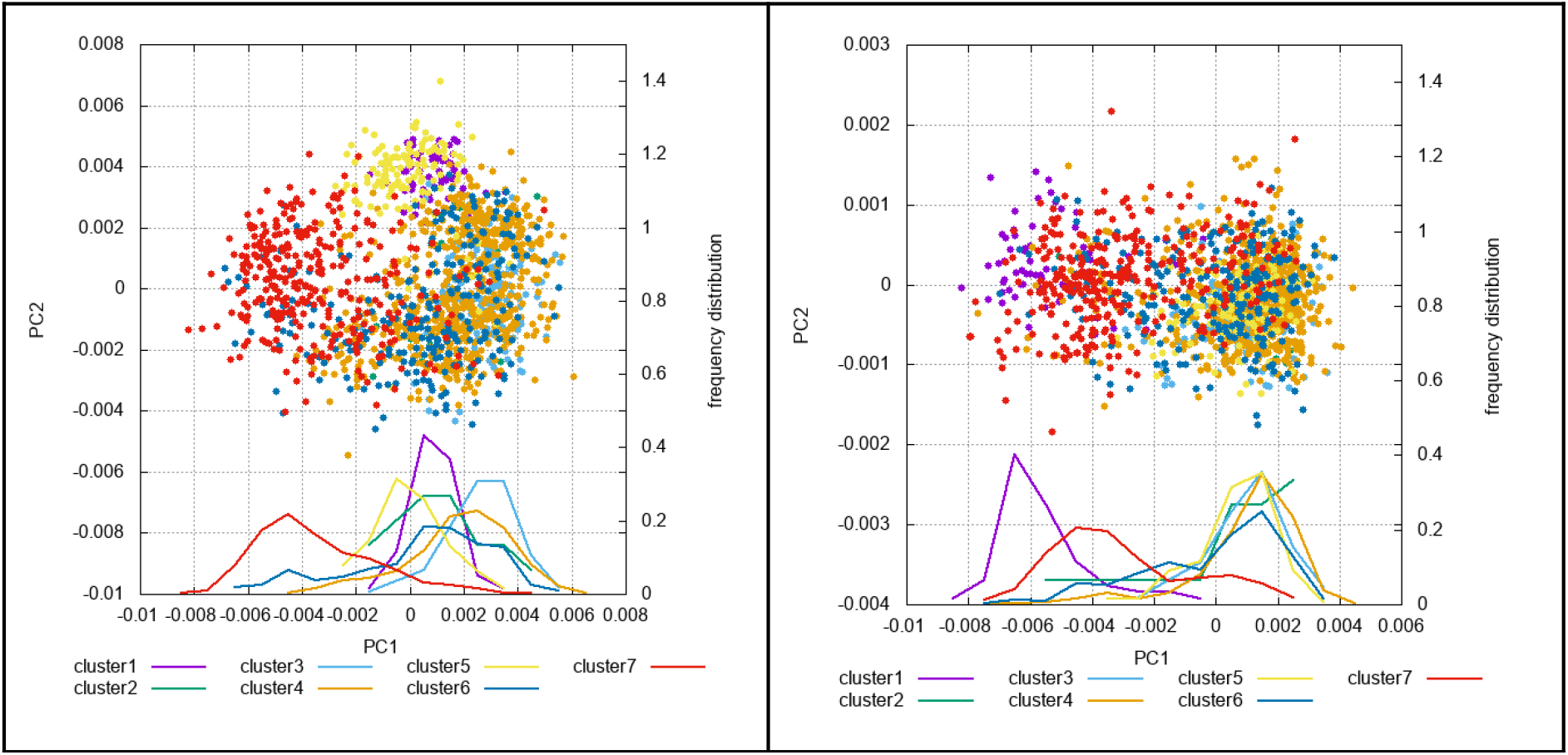
Interdomain clusters. **A:** plot of interdomain PC1 and PC2 of the clusters created from structures with tight interdomain interface. Experimental structures are included (FRET, open-like and closed-like are kindly provided by Mark Bowen) (Yanez Orozco et al., 2018). **B:** Representative structure of each cluster. **C:** Interface-forming sites in PDZ domain. **D:** Summary of the interdomain interface (first column), the complex-free form ratio (second column) and the ligand alignment (third column). **E:** Representative structure of cluster 5. **F**: Interdomain interaction (hydrogen-bonds and hydrophobic interaction, highlighted with blue and red, respectively) forming propensities of the residues in both domains, as found in cluster 5 (For interaction forming propensity, see Methods). **G-H:** Intradomain PC1 and PC2, with the clusters being colored, and their frequency distribution along PC1.

The orientation of the 4 C-terminal residues of the ligand depends on the interdomain torsional angle, therefore it also differs from cluster to cluster. The C-terminal tail of the ligand, pointing in the same direction, facilitates synergistic binding of the membrane receptors in the postsynaptic density, suggested and discussed by Zhang and coworkers (Long et al., 2003). However, in none of the tight orientations represented by these clusters are the two ligands orientated parallel to each other and in the same time perpendicular to the interdomain axis, which arrangement could allow for such synergistic binding. In many clusters, the perpendicular or antiparallel ligand orientation could allow one PDZ domain to bind a membrane receptor, and the other one to bind an intracellular partner. Nonetheless, it is not to be forgotten that all members of the clusters are composed of tight PDZ12 tandems, which only cover a fraction of the interdomain conformational space. Considering all the possible interdomain conformations, in which the two domains are not necessarily in interaction, there are many in which the ligands are in parallel orientation (Supplementary Figure 3).

Resulting from the interface-forming sites, the main interactions, which hold the two domains together, also differ from cluster to cluster. All interdomain hydrogen-bonds as well as hydrophobic interactions were determined in each conformation (for details, see Methods). Those interactions that are present in at least 10% of the conformations in any cluster are listed in Supplementary Table 2. For demonstration, the representative structure of cluster 5 and the residues involved in interdomain hydrogen bonds or hydrophobic interactions are shown on Figures 4.E and 4.F. Figure 4.F also shows that the interacting residues fall well within the previously defined interface-forming sites. The interdomain interface and the ligand orientation in the rest of the clusters is shown on Supplementary Figure 4.

PDZ12 tandem structures of PSD-95 determined previously by various experimental techniques were checked and compared to our clusters (Figure 4.A, Supplementary Figure 5). These experimental structures include, in particular, two chains of the same crystal structure from PDB (ID: 3gsl) (Sainlos et al., 2010), a structure determined by FRET presented by Bowen and coworkers (McCann et al., 2011) and two further structures, an open-like and a closed-like conformation, resulting from the refinement of the same FRET-structure by DMD simulations and presented by Sanabria and coworkers (Yanez Orozco et al., 2018) (5 structures in total). The FRET-structure falls outside of the circle of the interdomain conformational space covered by our simulations since it has a large interdomain bending angle due to the tail region of PDZ2 being inserted in between the two domains. The open-like and closed-like structures, however, fall within the covered interdomain conformational space. In the open-like conformation there are not many interdomain interactions, therefore it does not belong to any of the clusters composed exclusively by tight models. The closed-like conformation, on the other hand, lies at the edge of cluster 4, and, indeed, forms the interdomain interface between the sites termed ‘right side’ in both domains. It should be noted that were the two domains closer to one another in the open-like conformation, their relative orientation would allow for a larger interdomain interface between the right side of PDZ1 and the head of PDZ2, which would make it a member of cluster 3, rather than cluster 4 or 5, to which it is closer on the PCA plot. Also, both X-ray structures, lying on the edge of cluster 3, form the interdomain interface between the right side of PDZ1 and the head of PDZ2, which makes them indisputable members of cluster 3.

### Intra- and interdomain motions are interdependent

We analyzed the interdependence of intra- and interdomain motions. By plotting the principal components of intradomain motions and coloring each dot (representing a structure) according to the cluster of interdomain arrangement it belongs to, it is observed that intradomain conformations and interdomain orientations are not completely independent. (Figures 4.G and 4.H). It should be noted that the clusters contain both complexed and ligand-free structures. Furthermore, in PDZ2, cluster 1 and 7 are both slightly separated from the rest, even though complex forms are represented in a very high and low ratio in them, respectively.

As it was shown earlier, in PDZ1, the first principal component represents both the opening and closing of the binding site as well as the bending of β1-β2 and β2-β3 loops. In contrast, the binding site opening is suppressed by the bending of the β1-β2 loop in PDZ2. By comparing these regions to the sites introduced to define the interdomain interfaces, it is clear that β1-β2 constitutes the tail site while β2-β3 loop is involved in both the head region and right side (Supplementary Fig. 2). By looking at Figure 4.D, one can clearly see that in all but one clusters, at least one of these sites (i.e. the head or right side in PDZ1 or the tail in PDZ2) is involved in the interdomain interface.

Even though PDZ domains are traditionally categorized according only to the 4 C-terminal residues of their binding partner, the role of the upstream residues of the ligand has been emphasized recently (Luck et al., 2012). To gain further insight into the interaction between the PDZ domains and the ligand, hydrogen bonds were identified between both PDZ domains and the simulated cypin ligand, and their occurrences was calculated in percentages in all 7 clusters (Supplementary Table 3). Apart from 3 hydrogen-bonds present in high percentage in all clusters (between residues 0 and –2 in the ligand, and β2 strand in the PDZ domain), the rest of the bonds – many of them being formed between upstream ligand residues and the β2-β3 loop – show a different pattern in different clusters. This indicates a fine-tuned regulative interaction between ligand binding, intradomain dynamics and interdomain orientation.

Based on the above presented results, it is convenient to assume that interdomain orientation and the bound ligand are fine-tuning and regulating each other through a number of weak, atomic-level interactions. This might, in part, explain how the N-terminal PDZ domains of PSD-95 are capable of binding multimeric membrane-associated ligands as well as intracellular signaling proteins at the same time. This idea might fit well into the picture presented by Bowen and coworkers (McCann et al., 2012), according to which the PDZ12 tandem within the PSD-95 protein forms an independent conformational unit and is capable of performing detached motions from the other parts of the protein.

## Conclusion

The PSD-95 protein in the postsynaptic density of nerve cells performs numerous biological functions that require its peptide binding PDZ domains to be able to form versatile interactions with a number of binding partners. However, it is still unclear how PSD-95 discriminates between its ligands (McCann et al., 2011). The PDZ12 tandem of PSD-95 was shown to exhibit different function from the simple sum of the two costituting domains (Bach et al., 2012; Long et al., 2003; Wang et al., 2010), but the exact, atomic-level interactions between the two domains and the extent of interdomain reorientation remains elusive.

In this study, dynamic structural ensembles of the ligand-free and complexed forms of the PDZ12 tandem were created by externally restrained molecular dynamics simulations. ^1^H-^1^H NOEs were applied as distance restraints in both forms, and an additional ensemble was created of the complexed form in which S^2^ order parameter restraints were also incorporated. Applying the S^2^ restraints separately on each domain does not largely influence the interdomain conformational space, but it yields a description of the intradomain motions which is consistent with the experimental S^2^ parameters. Thus, we are capable of obtaining a description of possible interdomain orientations while keeping fast intradomain motions consistent with experimental observations. It is important to stress that no explicit restraining or other explicit influencing of interdomain interactions and orientation is present in our calculations. Restraining by S^2^ order parameters, representing motions on the fast ps-ns regime, is applied on the domains separately, allowing their independent, plausibly slower relative reorientation.

Our ensembles descriptive of the intra- and interdomain motions are in agreement with previous experimental findings, that is ligand binding changes the dynamics of the β2-β3 loop and it considerably expands the interdomain conformational space as well. The structural ensembles presented in this study provide a dynamic interpretation of the supramodular structural rearrangement of the PDZ12 tandem upon ligand binding. Our analysis is relevant in terms of the dynamics of the formation of the ligand-bound form and it gives an estimation on the redistribution of possible interdomain orientations of the PDZ12 tandem upon ligand binding and release.

The formation of the interdomain interface is not independent from the ligand binding. Our most important finding is that the different possible interdomain orientations, necessary to form certain interdomain interfaces, are influenced by the intradomain motions, or more specifically, by motions of the flexible and poorly conserved β2-β3 and β1-β2 loops in PDZ1, and of the β1-β2 loop in PDZ2 domain. The β2-β3 loop has already been suggested to play a key role in discriminating between different binding partners of PDZ domains (Luck et al., 2012), which is consistent with our analysis where we show that this loop is one of the regions most prone to forming interdomain interactions.

In summary, our results strongly suggest that ligand binding regulates interdomain orientation by modulating the probabilities of the formation of domain-domain interactions through different interdomain interfaces. It is also worth speculating that, vice versa, the relative orientation of the two domains in the PDZ12 tandem might have a role in shifting the preference for different binding partners, with the aid of the same β2-β3 loop. Additional – both in vitro and silico – investigation of the ligand-dependent interdomain dynamics of PDZ12 tandem of PSD-95 may shed light on its supramodular nature and help us further understand the molecular mechanisms behind the PSD-95 modulated rearrangement of the postsynaptic density in neuron cells.

## Methods

### Generation of structural ensembles

Dynamic structural ensembles were generated with externally restrained molecular dynamics simulation. In total, three structural ensembles were created, one of the free form, and two of the complex form of PDZ12 tandem. NOE restraints were applied to all three ensembles, while only one of the two complex ensembles was further restrained with S^2^ order parameters. Simulations were done by GROMACS 4.5.5 (Hess et al., 2008) with and in-house extension to handle NOE restraints in a pairwise manner over the replicas as well as to include S^2^ order parameter restraints as described previously (Fizil et al., 2015). The first (representative) model of the solution NMR structure of PDZ12 was used as input structure, deposited in the Protein Data Bank (PDB code: 2KA9) (Wang et al., 2009). For the free form, the two peptide ligands (chains B and C) were manually removed.

Experimental distance restraints were downloaded from the same PDB site in v2 format. Only ^1^H NOEs were used, in which stereospecific distance restraints were symmetrized and the resulting duplicates removed, which means 3169 ^1^H NOEs were retained of the original 3281. Experimental backbone S^2^ order parameters and chemical shifts (for N and H atoms) were kindly provided by Wenning Wang (Wang et al., 2009). In the simulations, 79 and 80 order parameters were used for the PDZ1 and PDZ2 domain, respectively. For data validation, 170 N and H chemical shifts were used. Distance and order parameter restraints were applied in the simulations according to the MUMO protocol (Best and Vendruscolo, 2004; Richter et al., 2007). For each system, 8 parallel simulations were carried out, in which NOEs were averaged in a pairwise manner, while order parameters were averaged for 8 parallel replicas.

To be able to describe the independent reorientation of the two PDZ domains, the S^2^ restraining routine was extended to handle local fitting for the calculation of S^2^ order parameters during the simulation independently for the two domains. In this implementation, a reference frame is assigned to each S^2^ parameter and the replicas are superimposed accordingly when calculating the instantaneous S^2^ parameters during each simulation step. In our particular case, when calculating S2 parameters for the first PDZ domain, the ensembles were superimposed on this domain and thus the S2 parameters were evaluated and compared to the experimental values in order to determine restraint energies and forces independently of the orientation of the second PDZ domain.

In the simulations, covalent bonds were constrained to retain their initial bond length by the LINCS algorithm. The simulations were carried out on 300 K, in TIP3P explicit solvent model. For each system, 8 parallel simulations were run for 20 ns simulation length, with 1 ps timestep. AMBER99SB-ILDN force field was applied. For temperature and pressure equilibration the first 2000 ps of each simulation was discarded. The final simulation trajectories were sampled with 100 ps frequency, which resulted in structural ensembles with 1448 replicas for each system.

### Analysis of structural ensembles

For validation of the structural ensembles, backbone RMSD values and experimental parameters were back-calculated from the ensembles (Table 1). Back-calculated chemical shifts were computed with shiftx2 (Han et al., 2011). Chemical shifts were than ensemble-averaged over all models in each ensembles and Pearson’s correlation was computed between the experimental and back-calculated chemical shifts. Back-calculated order parameters were computed with the CoNSEnsX+ webserver (Dudola et al., 2017). RMSD values were calculated for backbone atoms N, Cα2 helix. These motions mainly differ in the and C. Both order parameters and backbone RMSD were calculated separately for the two domains, after superimposing the structures on residues 4-88 and 99-186 for PDZ1 and PDZ2 domain, respectively. In case of the order parameters, back-calculated values for each domain were appended and their Pearson’s correlation determined against the experimental values.

Principal component analysis (PCA) and calculation of the overlap between various PCs was carried out by ProDy (Bakan et al., 2011, 2014). When determining intradomain motions, due to their large and in this context functionally irrelevant fluctuations, 3 residues from the flexible N- and C-terminal regions were excluded from the analysis, considering only the core region of both domains: residues 4-88 (PDZ1) and 96-186 (PDZ2). When determining interdomain motions, all structures were superimposed to a common PDZ1 core region template, and PCA was carried out for the core region of PDZ2 domain, without any further displacement. Structure figures were created with Chimera (Pettersen et al., 2004) and ChimeraX (Goddard et al., 2018), visual inspection of the internal motions was carried out by VMD (Humphrey et al., 1996).

### Identifying interdomain interactions

The size of the interdomain interface was calculated as the number of heavy atom-atom contacts between the core regions of the two domains within a cutoff range of 5 Å. Structures with less than 20 of such contacts were termed ‘loose’, while the rest were termed ‘tight’. In the three structural ensembles (1448 conformers each) there were 2722 loose and 1622 tight models in total. Interdomain distance was calculated as the distance between the geometric average of all Cα2 helix. These motions mainly differ in the atoms of the core region of the two domains.

Hydrogen bonds were identified based on angle-distance criteria (Baker and Hubbard, 1984; Xu et al., 1997). Interdomain hydrophobic interactions were identified as close contacts (within 5 Å cutoff range) between any carbon atom in the side chain of one of the following residues: Ala, Ile, Leu, Met, Phe, Pro, Thr, Trp, Val, Arg and Lys, or the Cα2 helix. These motions mainly differ in the atom of Gly. The interdomain interaction forming propensity of an amino acid residue in any cluster was calculated by counting the number of interdomain contacts (separately for hydrogen bonds and hydrophobic contacts) in which the given residue is involved, and dividing it by the number of models in the given cluster. This number might rise above 1.0 if a residue is involved in more than one hydrogen bonds or hydrophobic contacts.

### Calculating interdomain torsion angle

For the calculation of interdomain torsion angles, 16 data points were selected from the interdomain PCA plot (Figure 3.A-B), which are closest to 16 equidistant points lying on the best-fitting circle. Then, the best-fitting plane was determined to the centers of mass in the 3D space of the 16 different PDZ2 domain position. In the next step the centers of mass were projected on the 3D plane, and the center of the best-fitting circle was determined on the same plane. The 3D coordinates of the center of the circle in 3D space and of the normal vector of the plane (6 numbers in total) were calculated in the frame determined by the average position of Cα2 helix. These motions mainly differ in the atoms of Phe17, Ile57 and Val84. These residues were selected for being conserved, being robust in secondary structure (part of β2, β4 and β6, respectively) and having small RMSDs. To calculate the interdomain torsion angle of any PDZ12 tandem structure, the center of the 3D circle and its normal vector is determined relative to the Cα2 helix. These motions mainly differ in the atoms of the same three residues (Phe17, Ile57 and Val84) by using the coordinates obtained in the previous step.The Cα2 helix. These motions mainly differ in the of Phe17 and the center of mass of PDZ2 domain is projected on the plane, and the interdomain torsion angle results as the angle between these two projected points relative to the center of the 3D circle (Figure 3.D).

### Clustering of the ensembles

From the 3 generated ensembles, all tight models were selected (for details on the definition of tight models, see *Analysis of structural ensembles* in Methods). Based on the best-fitting circle fitted on the PC1 and PC2 coordinates of the interdomain principal components (as described in the previous section), their first and second principal components were converted into polar coordinates (*r* and *θ*). Using these polar coordinates, the tight models were clustered into 5 groups by complete linkage algorithm, and two of these clusters were further divided into 2 subclusters each. Outlier data points were removed after visual inspection. Representative structure of each cluster was identified as the one with the smallest RMSD from the average structure, computed for the backbone atoms (C, N and Cα) of the core region of PDZ2 domain.

## Acknowledgements

The authors acknowledge the support of the National Research, Development and Innovation Office – NKFIH through grants no. NF104198 and NN124363. Z.G. has been supported by a János Bolyai Research fellowship from the Hungarian Academy of Sciences. The research has been supported by the European Union, co-financed by the European Social Fund (EFOP-3.6.2-16-2017-00013, Thematic Fundamental Research Collaborations Grounding Innovation in Informatics and Infocommunications).

