## Supplementary Information for "Interdependence of intra- and inter-domain motions in the PSD-95 PDZ12 tandem"

Supplementary Figure 1.

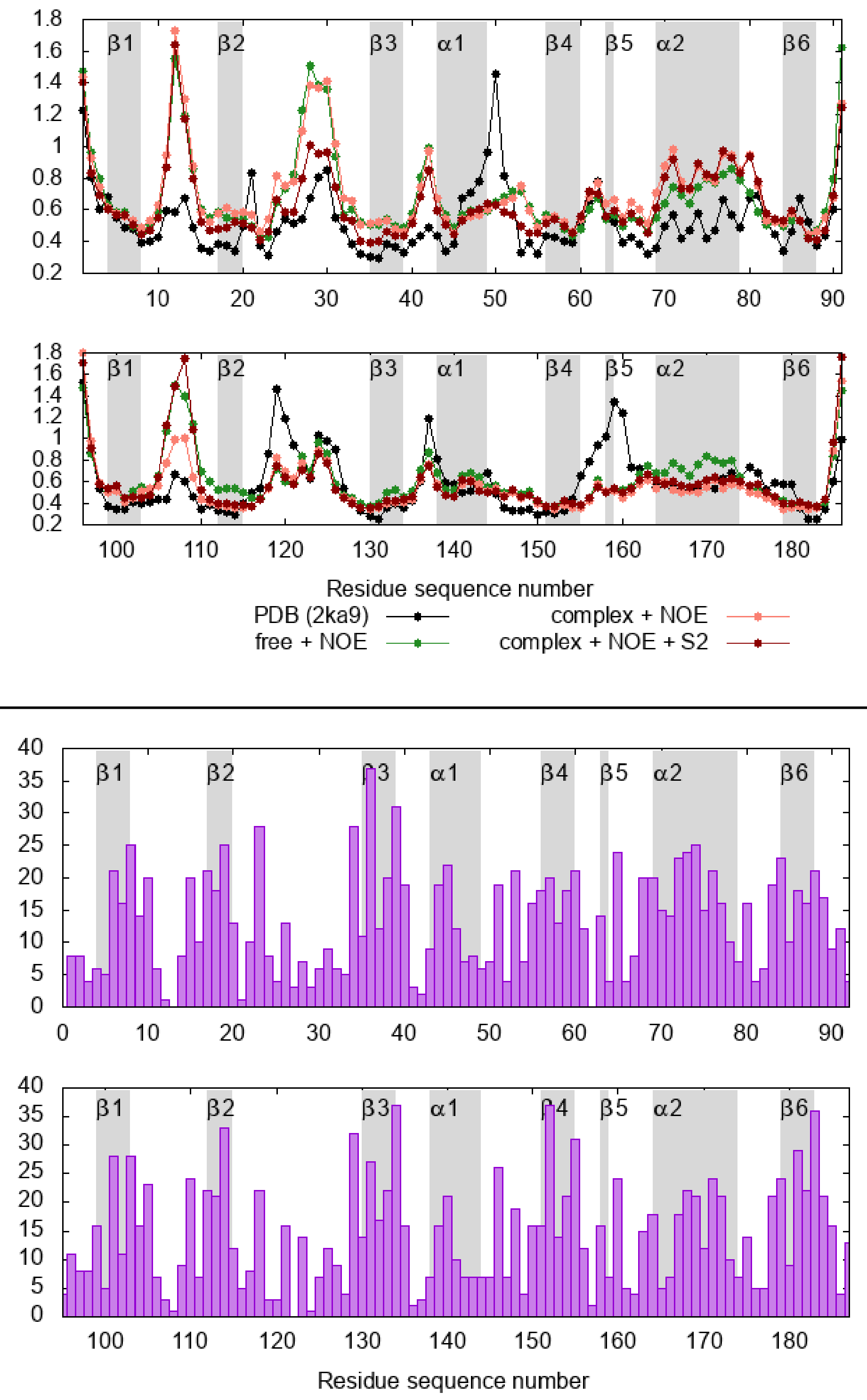

Local RMSDs of the original and generated ensembles (top) and the number of long-range NOE distance restraints per residue (bottom). NOEs were termed 'long-range' if they are restraining any two atoms between residues which are at least 2 position away from each other in the sequence.

Supplementary Figure 2.

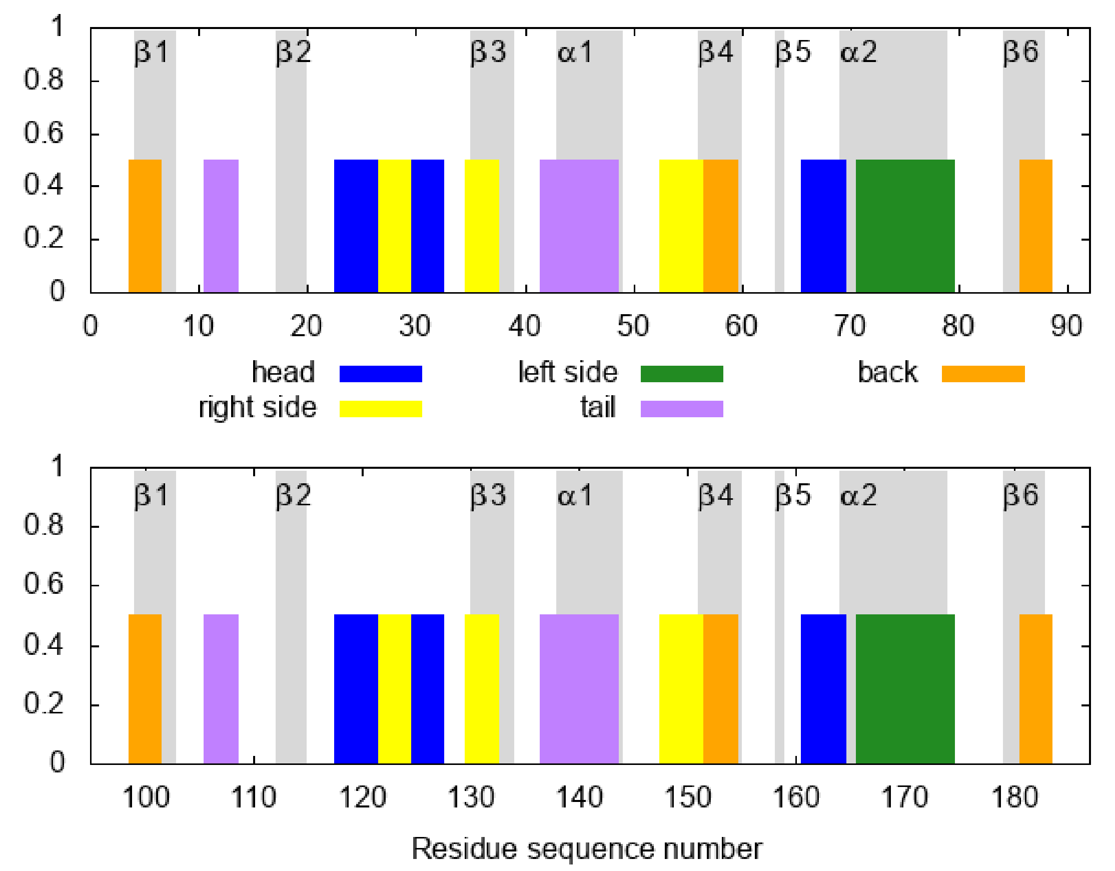

Sites in PDZ1 and PDZ2 domains. They are arbitrarily defined to facilitate identifying the interdomain interface. The color coding matches that of Figure 4.C in the main text.

Supplementary Table 1.

The 7 clusters created from the tight models, their size and composition, their interdomain interface, and the orientation and location of the ligand.

| Cluster | No. of Conformers | Complex form % | PDZ1 interface site | PDZ2 interface site | Ligand alignment (to each other) | Ligand alignment (to interdomain axis) | Ligand location |
|---|---|---|---|---|---|---|---|
| #1 | 60 | 100% | Head | Head | Antiparallel | Parallel | Opposite side |
| #2 | 15 | 100% | Head | Right side | Perpendicular | - | Same side |
| #3 | 152 | 100% | Right side | Head | Antiparallel | Parallel | Same side |
| #4 | 631 | 87% | Right side | Tail (Right side) | Antiparallel | Perpendicular | Same side |
| #5 | 131 | 100% | Right side | Back | Perpendicular | - | Opposite side |
| #6 | 250 | 70% | Tail | Head | Perpendicular | - | Same side |
| #7 | 317 | 4% | Tail | Tail | Perpendicular | - | Opposite side |

Supplementary Figure 3.

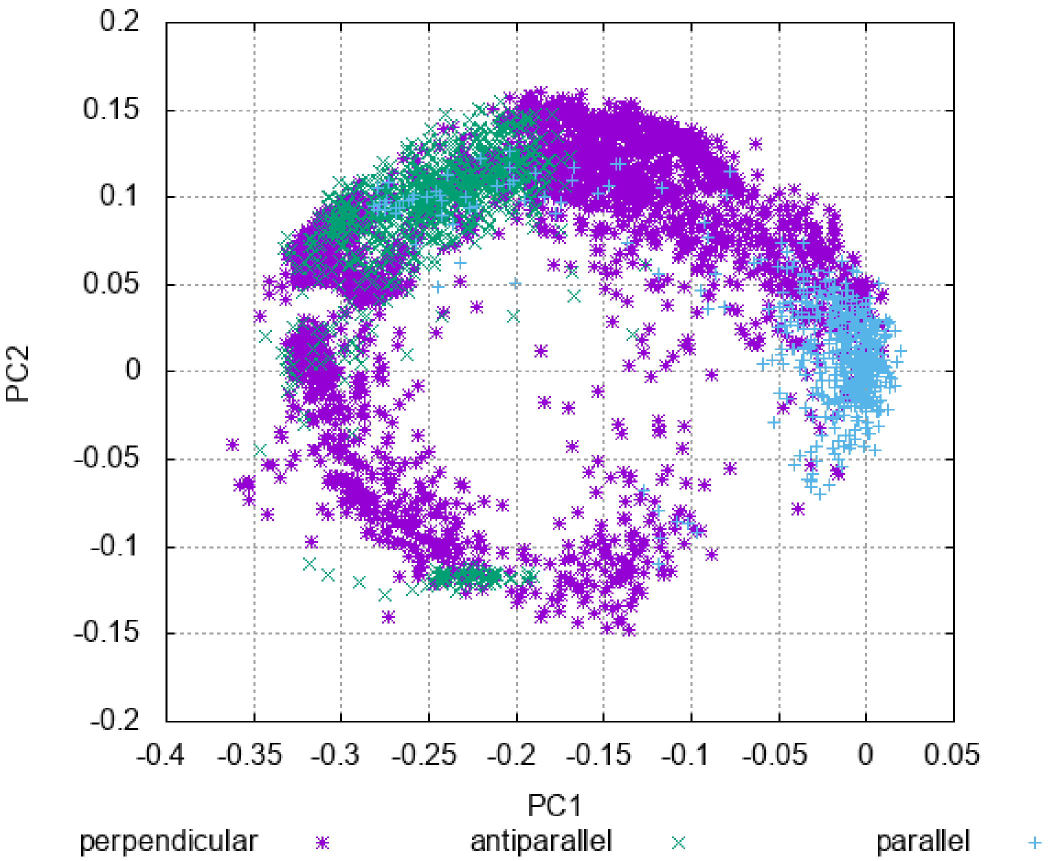

Distribution of the structures in the interdomain conformational space according to ligand orientation. The angle of the ligand was computed as the angle between the two vectors pointing from the CA atom of reside -2 to that of residue 0 of the ligands. The orientations were categorized according to the following: <45°: parallel, >45° and <135°: perpendicular, >135°: antiparallel.

Supplementary Table 2.

Occurrences of interdomain hydrogen bonds and hydrophobic interactions, with the two interacting atoms specified (rows) in every cluster (columns). Occurrences greater than or equal to 10% are highlighted with green background. Only those interactions are listed that occur in at least one of the clusters in a ratio greater than or equal to 10%.

| Hydrogen bonds | | | | | | | | |
|---|---|---|---|---|---|---|---|---|
| PDZ1 atom | PDZ2 atom | cl1 | cl2 | cl3 | cl4 | cl5 | cl6 | cl7 |
| ASN12@N | GLU136@OE2 | 0% | 0% | 0% | 0% | 0% | 0% | 12% |
| HIS27@ND1 | HIS141@NE2 | 0% | 0% | 0% | 26% | 0% | 0% | 0% |
| HIS27@NE2 | GLN121@OE1 | 0% | 0% | 26% | 0% | 0% | 0% | 0% |
| HIS27@NE2 | ILE123@O | 0% | 0% | 50% | 0% | 0% | 0% | 0% |
| HIS27@O | ARG145@NH1 | 0% | 0% | 0% | 38% | 0% | 0% | 0% |
| ASP30@OD1 | LYS133@NZ | 0% | 13% | 2% | 0% | 0% | 0% | 0% |
| ASP30@OD1 | LYS182@NZ | 20% | 0% | 0% | 0% | 0% | 0% | 0% |
| ASP30@OD2 | LYS133@NZ | 0% | 80% | 7% | 0% | 0% | 0% | 0% |
| ASP30@OD2 | LYS182@NZ | 15% | 0% | 0% | 0% | 0% | 0% | 0% |
| THR37@O | LYS182@NZ | 0% | 0% | 0% | 0% | 17% | 0% | 0% |
| LYS38@NZ | GLU100@OE1 | 0% | 0% | 0% | 1% | 45% | 0% | 0% |
| LYS38@NZ | GLU100@OE2 | 0% | 0% | 0% | 0% | 47% | 0% | 0% |
| LYS38@NZ | ILE134@O | 0% | 0% | 0% | 55% | 0% | 0% | 0% |
| LYS38@NZ | GLU136@OE1 | 0% | 0% | 0% | 39% | 0% | 0% | 0% |
| LYS38@NZ | GLU136@OE2 | 0% | 0% | 0% | 40% | 0% | 0% | 0% |
| GLY43@O | HIS141@NE2 | 0% | 0% | 0% | 0% | 0% | 8% | 19% |
| ALA46@O | HIS141@NE2 | 0% | 0% | 0% | 0% | 0% | 0% | 24% |
| GLN47@NE2 | ILE134@O | 0% | 0% | 0% | 0% | 0% | 6% | 51% |
| GLN47@O | ASN127@ND2 | 0% | 0% | 0% | 0% | 0% | 14% | 1% |
| GLN47@O | ILE148@N | 0% | 0% | 0% | 0% | 0% | 1% | 16% |
| ASP48@O | ILE148@N | 0% | 0% | 0% | 0% | 0% | 0% | 16% |
| ASP48@OD2 | ILE134@N | 0% | 0% | 0% | 0% | 0% | 0% | 12% |
| ARG52@NE | PRO124@O | 0% | 0% | 0% | 0% | 0% | 11% | 0% |
| ARG52@NH1 | GLN147@OE1 | 0% | 0% | 0% | 0% | 0% | 2% | 19% |
| ARG52@NH2 | GLN147@OE1 | 0% | 0% | 0% | 1% | 0% | 4% | 34% |
| ASN54@ND2 | GLN147@OE1 | 0% | 0% | 0% | 25% | 0% | 0% | 0% |
| ARG66@NH1 | GLU161@OE1 | 45% | 0% | 0% | 0% | 0% | 0% | 0% |
| ARG66@NH1 | GLU161@OE2 | 32% | 0% | 0% | 0% | 0% | 0% | 0% |
| ARG66@NH2 | GLU161@OE1 | 32% | 0% | 0% | 0% | 0% | 0% | 0% |
| ARG66@NH2 | GLU161@OE2 | 35% | 0% | 0% | 0% | 0% | 0% | 0% |
| TYR87@OH | PRO124@O | 17% | 0% | 0% | 0% | 0% | 0% | 0% |
| GLU9@OE1 | LYS133@NZ | 0% | 0% | 0% | 0% | 0% | 0% | 32% |
| GLU9@OE2 | LYS133@NZ | 0% | 0% | 0% | 0% | 0% | 0% | 24% |
| Hydrophobic interactions | | | | | | | | |
| PDZ1 atom | PDZ2 atom | cl1 | cl2 | cl3 | cl4 | cl5 | cl6 | cl7 |
| ILE28@CB | ILE148@CG2 | 0% | 13% | 1% | 0% | 0% | 0% | 0% |

| | | | | | | | | |
|---|---|---|---|---|---|---|---|---|
| ILE28@CG1 | ILE148@CG2 | 0% | 13% | 3% | 0% | 0% | 0% | 0% |
| ILE28@CG2 | ILE148@CG2 | 0% | 13% | 5% | 0% | 0% | 0% | 0% |
| PHE35@CB | PRO124@CB | 0% | 0% | 86% | 0% | 0% | 0% | 0% |
| PHE35@CB | PRO124@CG | 0% | 0% | 74% | 0% | 0% | 0% | 0% |
| PHE35@CD1 | PRO124@CB | 0% | 0% | 80% | 0% | 0% | 0% | 0% |
| PHE35@CD1 | PRO124@CD | 0% | 0% | 18% | 0% | 0% | 0% | 0% |
| PHE35@CD1 | PRO124@CG | 0% | 0% | 87% | 0% | 0% | 0% | 0% |
| PHE35@CD2 | PRO124@CB | 0% | 0% | 63% | 0% | 0% | 0% | 0% |
| PHE35@CD2 | PRO124@CD | 0% | 0% | 18% | 0% | 0% | 0% | 0% |
| PHE35@CD2 | PRO124@CG | 0% | 0% | 78% | 0% | 0% | 0% | 0% |
| PHE35@CE1 | PRO124@CB | 0% | 0% | 49% | 0% | 0% | 0% | 0% |
| PHE35@CE1 | PRO124@CD | 0% | 0% | 17% | 0% | 0% | 0% | 0% |
| PHE35@CE1 | PRO124@CG | 0% | 0% | 81% | 0% | 0% | 0% | 0% |
| PHE35@CE2 | PRO124@CB | 0% | 0% | 34% | 0% | 0% | 0% | 0% |
| PHE35@CE2 | PRO124@CD | 0% | 0% | 13% | 0% | 0% | 0% | 0% |
| PHE35@CE2 | PRO124@CG | 0% | 0% | 64% | 0% | 0% | 0% | 0% |
| PHE35@CG | PRO124@CB | 0% | 0% | 84% | 0% | 0% | 0% | 0% |
| PHE35@CG | PRO124@CD | 0% | 0% | 20% | 0% | 0% | 0% | 0% |
| PHE35@CG | PRO124@CG | 0% | 0% | 86% | 0% | 0% | 0% | 0% |
| PHE35@CZ | PRO124@CB | 0% | 0% | 28% | 0% | 0% | 0% | 0% |
| PHE35@CZ | PRO124@CD | 0% | 0% | 16% | 0% | 0% | 0% | 0% |
| PHE35@CZ | PRO124@CG | 0% | 0% | 64% | 0% | 0% | 0% | 0% |
| THR37@CG2 | PRO124@CB | 0% | 0% | 41% | 0% | 0% | 0% | 0% |
| THR37@CG2 | GLY125@CA | 0% | 0% | 74% | 1% | 0% | 0% | 0% |
| THR37@CG2 | ILE134@CB | 0% | 0% | 0% | 17% | 0% | 0% | 0% |
| THR37@CG2 | ILE134@CG1 | 0% | 0% | 0% | 12% | 0% | 0% | 0% |
| THR37@CG2 | ILE134@CG2 | 0% | 0% | 0% | 22% | 0% | 0% | 0% |
| LYS38@CB | ILE134@CB | 0% | 0% | 0% | 14% | 0% | 0% | 0% |
| LYS38@CB | ILE148@CB | 0% | 0% | 0% | 23% | 0% | 0% | 0% |
| LYS38@CB | ILE148@CG1 | 0% | 0% | 0% | 61% | 0% | 0% | 0% |
| LYS38@CB | LYS182@CD | 0% | 0% | 0% | 0% | 26% | 0% | 0% |
| LYS38@CD | LYS133@CG | 0% | 0% | 0% | 18% | 0% | 0% | 0% |
| LYS38@CD | ILE134@CB | 0% | 0% | 0% | 39% | 0% | 0% | 0% |
| LYS38@CD | ILE134@CG2 | 0% | 0% | 0% | 24% | 0% | 0% | 0% |
| LYS38@CD | ILE148@CG1 | 0% | 0% | 0% | 14% | 0% | 0% | 0% |
| LYS38@CG | ILE134@CB | 0% | 0% | 0% | 43% | 0% | 0% | 0% |
| LYS38@CG | ILE134@CG1 | 0% | 0% | 0% | 21% | 0% | 0% | 0% |
| LYS38@CG | ILE134@CG2 | 0% | 0% | 0% | 26% | 0% | 0% | 0% |
| LYS38@CG | ILE148@CG1 | 0% | 0% | 0% | 27% | 0% | 0% | 0% |
| LYS38@CG | LYS182@CD | 0% | 0% | 0% | 0% | 21% | 0% | 0% |
| ILE39@CB | PRO124@CB | 0% | 0% | 0% | 0% | 0% | 41% | 0% |
| ILE39@CB | PRO124@CG | 0% | 0% | 0% | 0% | 0% | 23% | 0% |
| ILE39@CG1 | PRO124@CB | 0% | 0% | 0% | 0% | 0% | 44% | 0% |
| ILE39@CG1 | PRO124@CD | 0% | 0% | 0% | 0% | 0% | 11% | 0% |
| ILE39@CG1 | PRO124@CG | 0% | 0% | 0% | 0% | 0% | 25% | 0% |
| ILE39@CG2 | PRO124@CB | 0% | 0% | 0% | 0% | 0% | 31% | 1% |

| | | | | | | | | |
|---|---|---|---|---|---|---|---|---|
| ILE39@CG2 | PRO124@CG | 0% | 0% | 0% | 0% | 0% | 14% | 0% |
| PRO41@CB | GLY119@CA | 0% | 0% | 0% | 0% | 0% | 13% | 0% |
| ALA46@CB | PRO124@CB | 0% | 0% | 0% | 0% | 0% | 25% | 1% |
| ALA46@CB | GLY125@CA | 0% | 0% | 0% | 0% | 0% | 11% | 0% |
| GLY49@CA | PRO124@CB | 0% | 0% | 0% | 0% | 0% | 26% | 1% |
| GLY49@CA | GLY125@CA | 0% | 0% | 0% | 0% | 0% | 70% | 2% |
| ARG50@CB | ILE148@CB | 0% | 0% | 0% | 0% | 0% | 0% | 11% |
| ARG52@CB | PRO124@CB | 0% | 0% | 0% | 0% | 0% | 16% | 0% |
| ARG52@CB | PRO124@CG | 0% | 0% | 0% | 0% | 0% | 21% | 0% |
| ARG52@CG | PRO124@CB | 0% | 0% | 0% | 0% | 0% | 11% | 0% |
| ARG52@CG | PRO124@CG | 0% | 0% | 0% | 0% | 0% | 18% | 0% |
| VAL53@CB | GLY125@CA | 0% | 0% | 35% | 0% | 0% | 0% | 0% |
| VAL53@CB | LEU153@CD1 | 0% | 0% | 0% | 0% | 22% | 0% | 0% |
| VAL53@CG1 | PRO124@CB | 0% | 0% | 1% | 0% | 0% | 24% | 0% |
| VAL53@CG1 | PRO124@CG | 0% | 0% | 1% | 0% | 0% | 15% | 0% |
| VAL53@CG1 | LEU153@CD1 | 0% | 0% | 0% | 0% | 43% | 0% | 0% |
| VAL53@CG1 | LEU153@CD2 | 0% | 0% | 0% | 0% | 11% | 0% | 0% |
| VAL53@CG1 | LYS182@CD | 0% | 0% | 0% | 0% | 18% | 0% | 0% |
| VAL53@CG2 | PRO124@CB | 0% | 0% | 26% | 0% | 0% | 8% | 0% |
| VAL53@CG2 | GLY125@CA | 0% | 0% | 78% | 0% | 0% | 0% | 0% |
| VAL53@CG2 | ILE148@CB | 0% | 0% | 0% | 14% | 0% | 0% | 0% |
| VAL53@CG2 | LEU153@CD1 | 0% | 0% | 0% | 0% | 92% | 0% | 0% |
| VAL53@CG2 | LEU153@CD2 | 0% | 0% | 0% | 0% | 14% | 0% | 0% |
| VAL53@CG2 | LEU153@CG | 0% | 0% | 0% | 0% | 15% | 0% | 0% |
| VAL53@CG2 | LYS182@CD | 0% | 0% | 0% | 0% | 27% | 0% | 0% |
| PHE59@CD2 | GLY125@CA | 15% | 0% | 0% | 0% | 0% | 0% | 0% |
| PHE59@CE1 | GLY125@CA | 15% | 0% | 0% | 0% | 0% | 0% | 0% |
| PHE59@CE2 | GLY125@CA | 32% | 0% | 0% | 0% | 0% | 0% | 0% |
| PHE59@CZ | GLY125@CA | 33% | 0% | 0% | 0% | 0% | 0% | 0% |

Supplementary Figure 4.

Supplementary figures about each cluster. Top: interdomain interaction forming propensity of all residues in both domains. The hydrogen bonds and the hydrophobic interactions are highlighted with blue and red, respectively. For the calculation of interdomain interaction forming propensity, see Methods. Middle: the representative structure of the given cluster, with the interdomain interactions highlighted, hydrogen bonds with blue, hydrophobic interactions with red. Bottom: the representative structure of the given cluster, with the 4 C-terminal residues of the peptide ligand shown in red. The orientation of the middle and bottom figures might differ to give better visibility on the interactions and the ligand orientations, respectively. In all figures, PDZ1 is on the left side and PDZ2 is on the right side.

<table>
<tr><th>Cluster 1</th><th>Cluster 2</th></tr>
<tr><td>
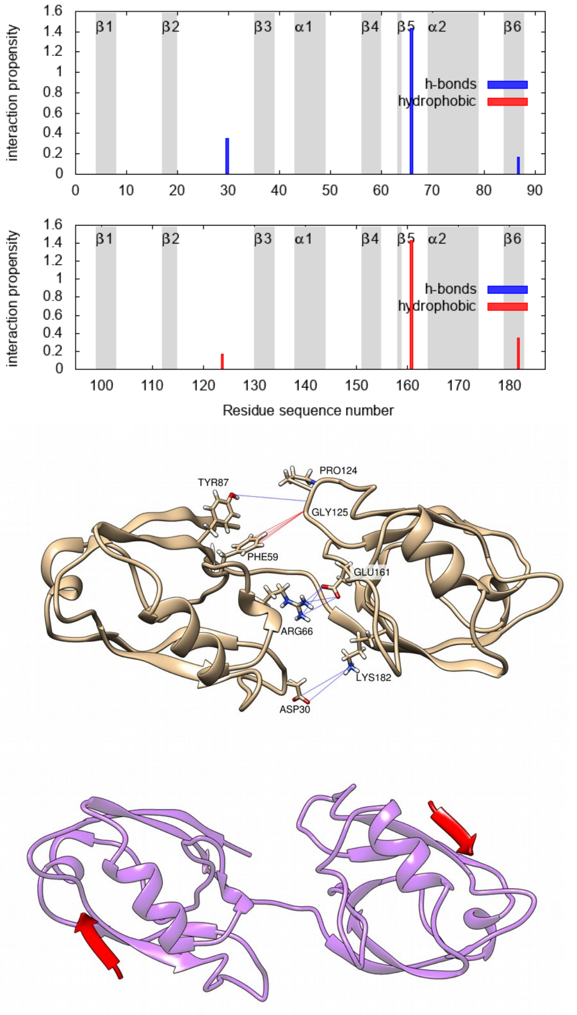

</td><td>
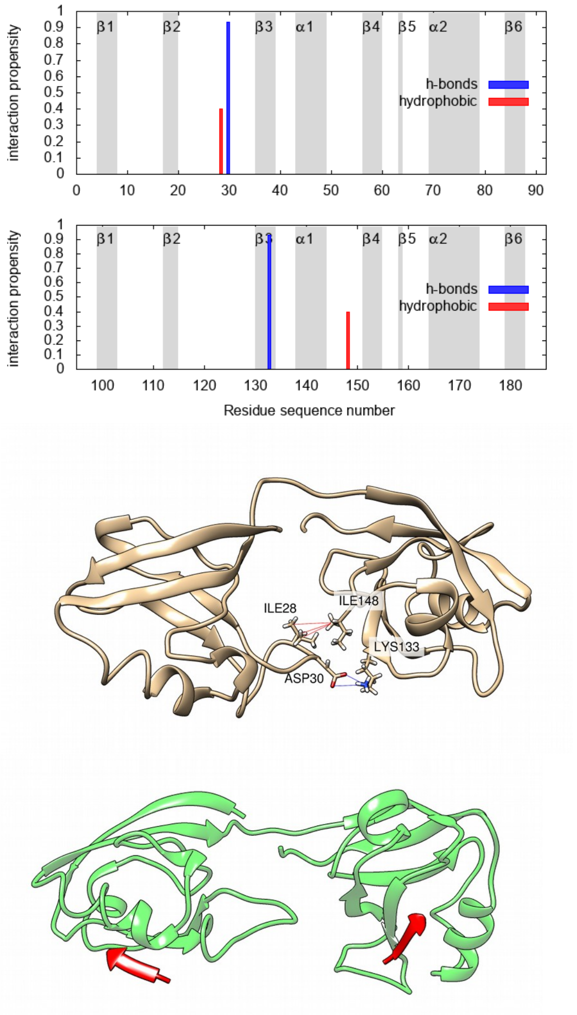

</td></tr>
<tr><th>Cluster 3</th><th>Cluster 4</th></tr>
</table>

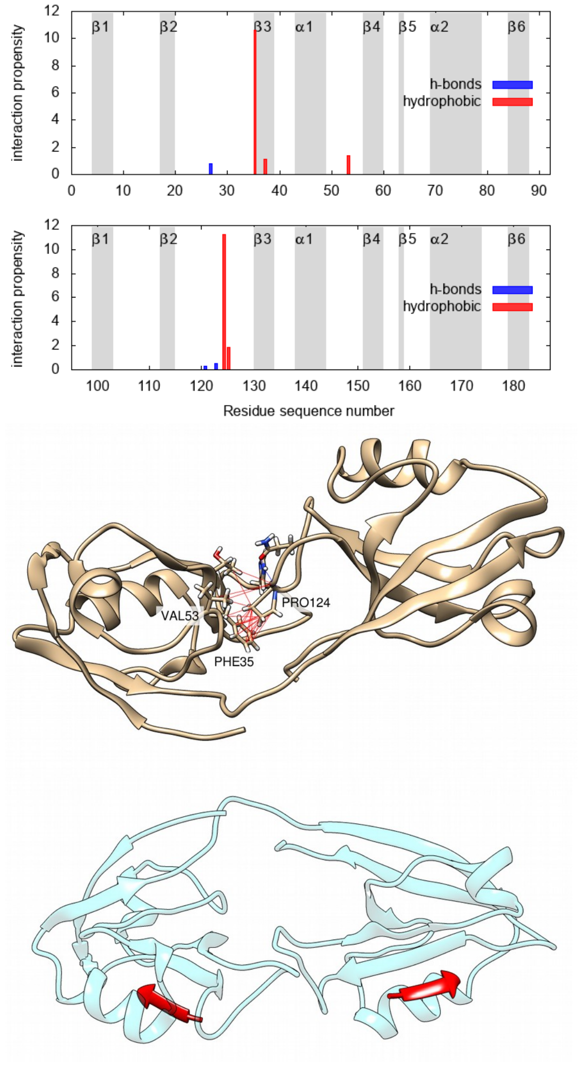

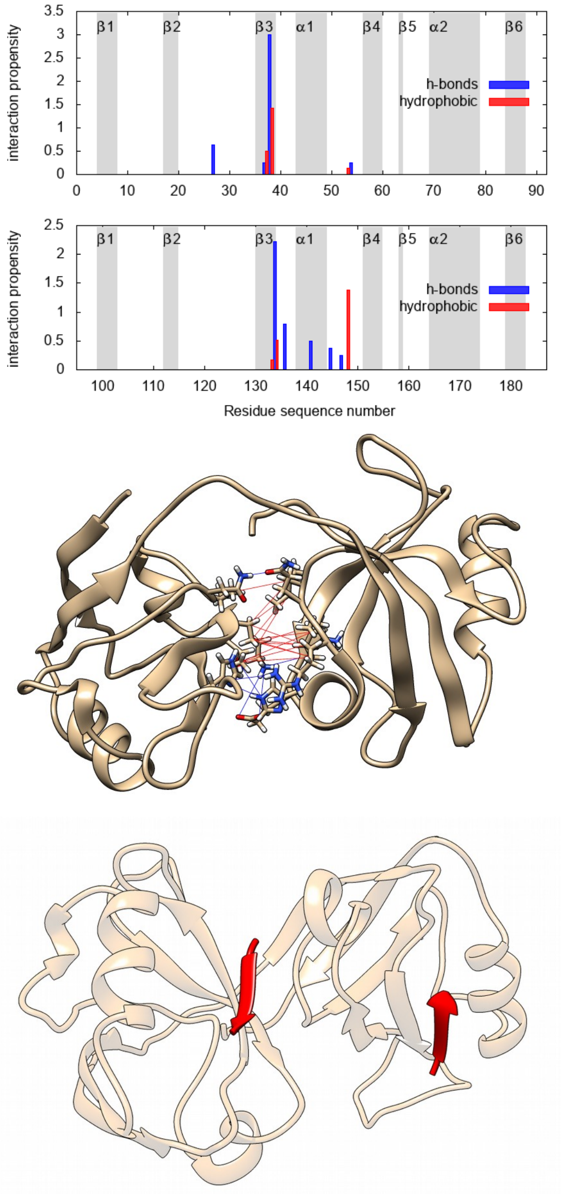

### Cluster 5

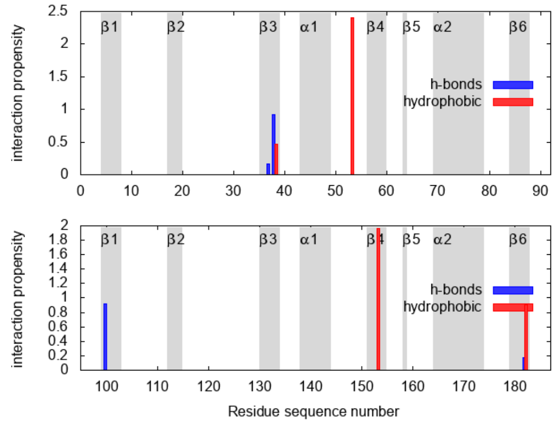

### Cluster 6

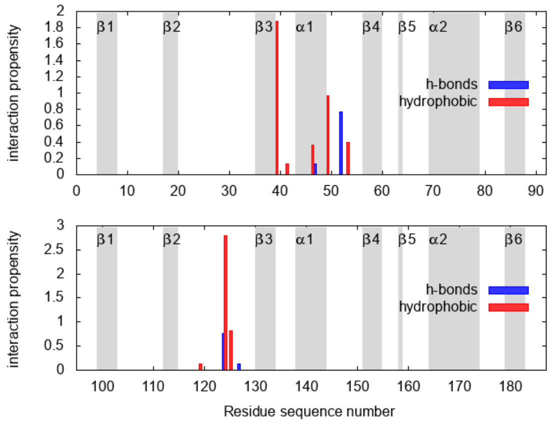

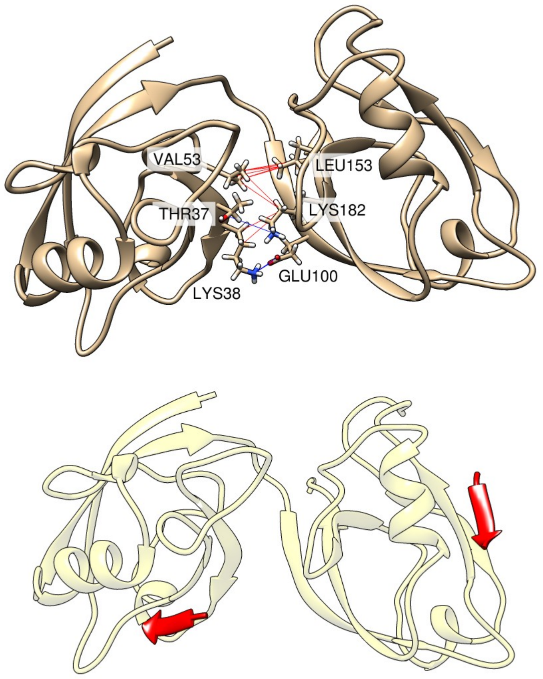
VAL53
LEU153
THR37
LYS182
LYS38
GLU100

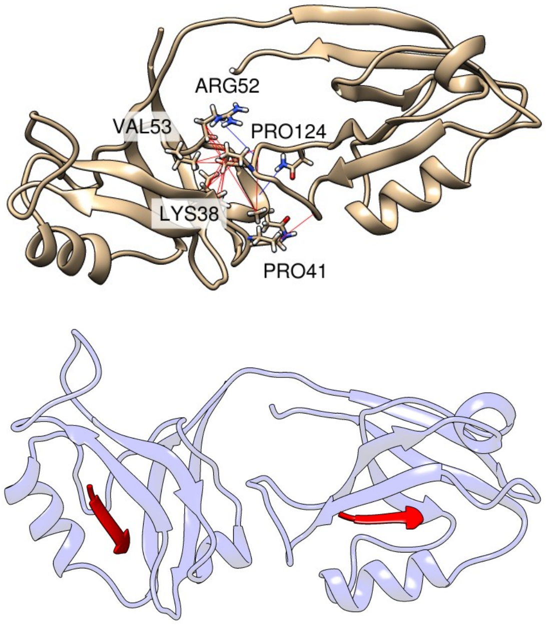
ARG52
VAL53
PRO124
LYS38
PRO41

### Cluster 7

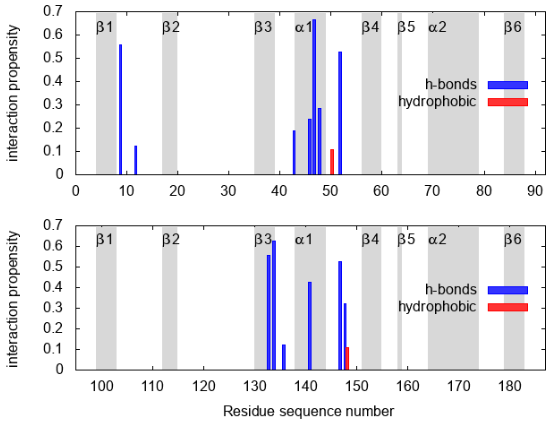
interaction propensity
β1
β2
β3
α1
β4
β5
α2
β6
h-bonds
hydrophobic
0
0.1
0.2
0.3
0.4
0.5
0.6
0.7
0 10 20 30 40 50 60 70 80 90
100 110 120 130 140 150 160 170 180
Residue sequence number

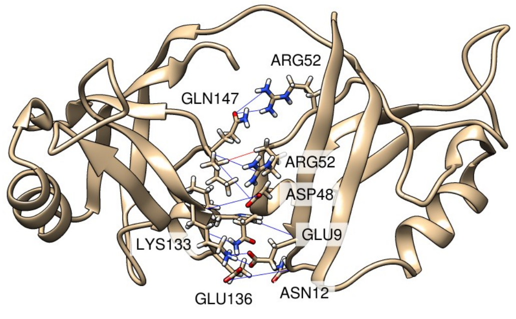
ARG52
GLN147
ARG52
ASP48
GLU9
LYS133
GLU136
ASN12

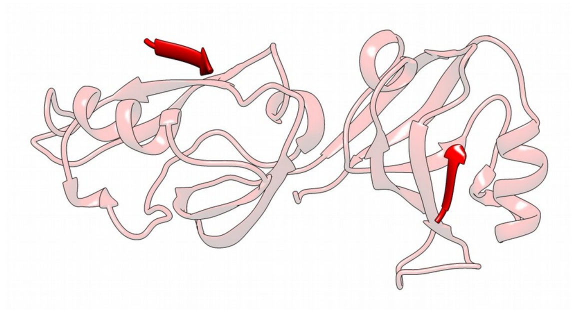

Supplementary Table 3.

Hydrogen bonds between the domains and the ligands. For every hydrogen bond, one atom from the PDZ domain and one from the ligand is specified. Three hydrogen bonds occurring in very frequently in both domains are highlighted with green. Occurrences greater than or equal to 10% are highlighted with light blue. Only those interactions are listed that occur in at least one of the clusters in a ratio greater than or equal to 10%.

| Almost always occurring PDZ-ligand hydrogen bonds (≥50% in all) | | | | | | | | | | | |
|---|---|---|---|---|---|---|---|---|---|---|---|
| Frequentyl occurring PDZ-ligand hydrogen bonds (≥10%) | | | | | | | | | | | |
| **PDZ1** | | | | | | | | | | | |
| PDZ atom | ligand atom | PDZ region | ligand residue | all | cl1 | cl2 | cl3 | cl4 | cl5 | cl6 | cl7 |
| PHE17@O | VAL198@N | β2 | 0 | 83% | 87% | 80% | 89% | 82% | 78% | 84% | 79% |
| ILE19@N | SER196@O | β2 | -2 | 98% | 100% | 100% | 99% | 98% | 96% | 98% | 100% |
| ILE19@O | SER195@OG | β2 | -3 | 15% | 18% | 0% | 12% | 16% | 15% | 18% | 36% |
| ILE19@O | SER196@N | β2 | -2 | 94% | 100% | 100% | 93% | 91% | 99% | 94% | 100% |
| GLY21@N | GLN190@O | β2 | -8 | 4% | 0% | 73% | 10% | 1% | 0% | 0% | 0% |
| GLY21@N | PHE194@O | β2 | -4 | 22% | 12% | 0% | 3% | 40% | 3% | 9% | 36% |
| ASP24@O | VAL191@N | β2-β3 | -7 | 1% | 0% | 0% | 0% | 0% | 0% | 2% | 21% |
| ASN25@ND2 | VAL192@O | β2-β3 | -6 | 21% | 0% | 0% | 1% | 42% | 0% | 2% | 0% |
| ASN25@ND2 | PRO193@O | β2-β3 | -5 | 1% | 17% | 0% | 0% | 0% | 0% | 0% | 0% |
| ASN25@OD1 | VAL191@N | β2-β3 | -7 | 3% | 0% | 13% | 1% | 0% | 0% | 11% | 29% |
| ASN25@OD1 | VAL192@N | β2-β3 | -6 | 2% | 0% | 0% | 1% | 2% | 0% | 5% | 50% |
| HIS27@ND1 | VAL192@N | β2-β3 | -6 | 1% | 0% | 20% | 5% | 0% | 0% | 0% | 0% |
| HIS27@O | GLN190@NE2 | β2-β3 | -8 | 2% | 0% | 0% | 11% | 0% | 2% | 1% | 0% |
| HIS27@O | VAL191@N | β2-β3 | -7 | 8% | 0% | 0% | 0% | 0% | 63% | 1% | 0% |
| HIS27@O | VAL192@N | β2-β3 | -6 | 3% | 0% | 7% | 0% | 0% | 6% | 14% | 0% |
| THR37@OG1 | SER195@OG | β3 | -3 | 14% | 2% | 67% | 19% | 2% | 46% | 17% | 29% |
| HIS70@NE2 | VAL192@O | α2 | -6 | 4% | 42% | 0% | 0% | 3% | 0% | 0% | 0% |
| HIS70@NE2 | SER196@OG | α2 | -2 | 7% | 5% | 33% | 14% | 7% | 5% | 5% | 0% |
| **PDZ2** | | | | | | | | | | | |
| PDZ atom | ligand atom | | | all | cl1 | cl2 | cl3 | cl4 | cl5 | cl6 | cl7 |
| PHE112@O | SER206@OG | β2 | -1 | 14% | 5% | 27% | 13% | 16% | 18% | 9% | 7% |
| PHE112@O | VAL207@N | β2 | 0 | 53% | 88% | 60% | 59% | 47% | 57% | 52% | 43% |
| SER113@OG | SER204@O | β2 | -3 | 1% | 0% | 13% | 2% | 0% | 0% | 1% | 0% |
| SER113@OG | SER205@O | β2 | -2 | 14% | 2% | 20% | 11% | 15% | 12% | 14% | 21% |
| SER113@OG | SER206@OG | β2 | -1 | 8% | 28% | 20% | 5% | 5% | 9% | 10% | 7% |
| ILE114@N | SER205@O | β2 | -2 | 95% | 98% | 87% | 93% | 95% | 98% | 94% | 100% |
| ILE114@O | SER204@OG | β2 | -3 | 16% | 20% | 7% | 11% | 19% | 18% | 12% | 7% |
| ILE114@O | SER205@N | β2 | -2 | 76% | 83% | 40% | 80% | 77% | 86% | 70% | 50% |
| ASN120@ND2 | VAL201@O | β2-β3 | -6 | 6% | 0% | 0% | 0% | 11% | 5% | 0% | 0% |

| | | | | | | | | | | | |
|---|---|---|---|---|---|---|---|---|---|---|---|
| ASN120@ND2 | PRO202@O | β2-β3 | -5 | 5% | 5% | 0% | 0% | 1% | 34% | 0% | 0% |
| ASN120@ND2 | PHE203@O | β2-β3 | -4 | 8% | 62% | 0% | 14% | 3% | 12% | 1% | 0% |
| ASN120@OD1 | VAL201@N | β2-β3 | -6 | 6% | 0% | 0% | 0% | 7% | 2% | 13% | 0% |
| ASN120@OD1 | PHE203@N | β2-β3 | -4 | 1% | 20% | 0% | 0% | 0% | 1% | 0% | 0% |
| HIS122@NE2 | VAL201@N | β2-β3 | -6 | 1% | 0% | 0% | 0% | 0% | 1% | 5% | 14% |
| HIS165@NE2 | PRO202@O | α2 | -5 | 1% | 12% | 0% | 0% | 0% | 2% | 0% | 0% |
| HIS165@NE2 | SER205@OG | α2 | -2 | 5% | 48% | 0% | 13% | 1% | 0% | 2% | 0% |
| GLU166@OE1 | VAL200@N | α2 | -7 | 1% | 0% | 13% | 3% | 0% | 0% | 0% | 0% |
| GLU166@OE1 | VAL201@N | α2 | -6 | 1% | 0% | 13% | 3% | 0% | 0% | 0% | 0% |
| GLU166@OE2 | GLN199@NE2 | α2 | -8 | 2% | 3% | 0% | 0% | 0% | 10% | 3% | 14% |
| GLU166@OE2 | VAL200@N | α2 | -7 | 1% | 0% | 40% | 2% | 0% | 0% | 0% | 0% |
| GLU166@OE2 | VAL201@N | α2 | -6 | 1% | 0% | 33% | 2% | 0% | 0% | 0% | 0% |
| LYS173@NZ | SER206@O | α2 | -1 | 5% | 0% | 7% | 6% | 5% | 5% | 3% | 14% |

Supplementary Figure 5.

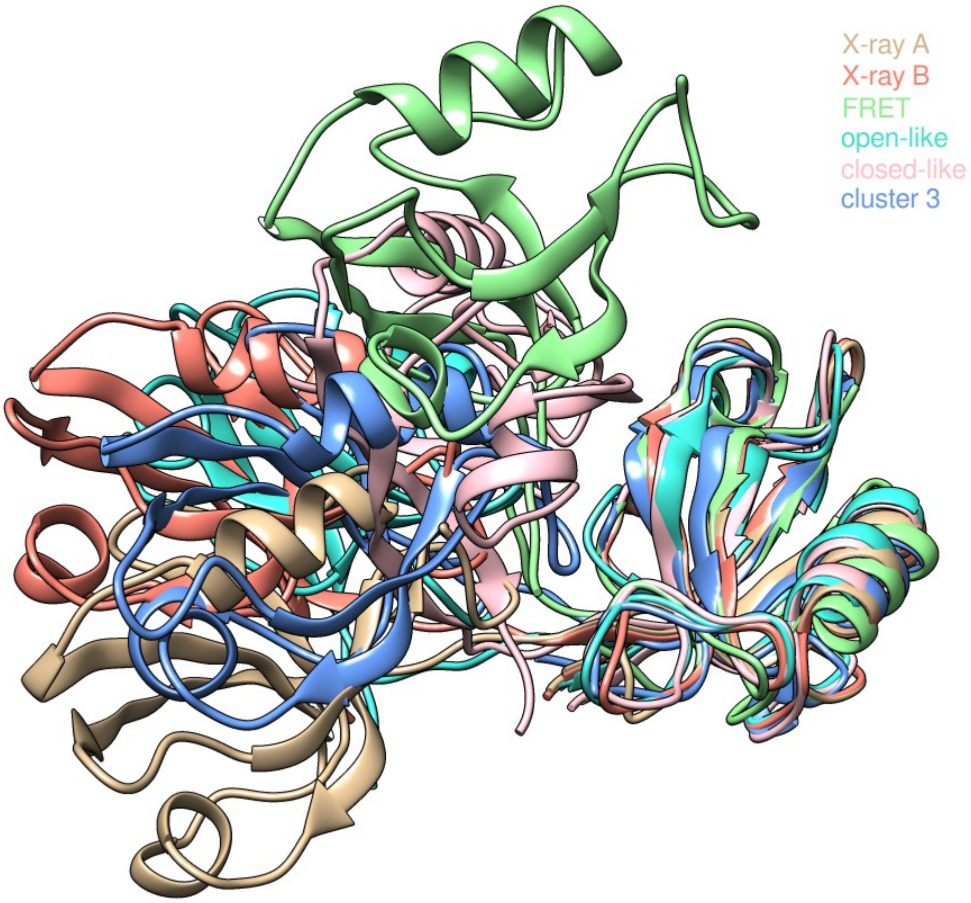

Structures of PSD-95 PDZ12 tandem determined by various experimental methods. The structures are superimposed on a common PDZ1 template so the difference in interdomain orientation becomes apparent by the displacement of the PDZ2 domain. X-ray structures are obtained from PDB (PDB code: 3GSL) (Sainlos et al., 2010), the FRET, open-like and closed-like structures were kindly provided by Mark Bowen (McCann et al., 2011, Yanez Orozco et al., 2018), and the representative structure of cluster 3 is the result of the presented work.

Sainlos, M., Tigaret, C., Poujol, C., Olivier, N.B., Bard, L., Breillat, C., Thiolon, K., Choquet, D., and Imperiali, B. (2010). Biomimetic divalent ligands for the acute disruption of synaptic AMPAR stabilization. Nat. Chem. Biol. *7*, 81.

McCann, J.J.J., Zheng, L., Chiantia, S., and Bowen, M.E.E. (2011). Domain Orientation in the N-Terminal PDZ Tandem from PSD-95 Is Maintained in the Full-Length Protein. Structure *19*, 810–820.

Yanez Orozco, I.S., Mindlin, F.A., Ma, J., Wang, B., Levesque, B., Spencer, M., Rezaei Adariani, S., Hamilton, G., Ding, F., Bowen, M.E., et al. (2018). Identifying weak interdomain interactions that stabilize the supertertiary structure of the N-terminal tandem PDZ domains of PSD-95. Nat. Commun. *9*, 3724.
